# Metrics matter: Modelling plant-frugivore networks in a tropical dry forest ecosystem

**DOI:** 10.64898/2026.09.28.755107

**Authors:** Allegra N. DePasquale, Theo Batchelor, Blanca Cejalvo Insausti, Jorja Strickland, Natalie Sanchez, Daniel Mennill, Omer Nevo, Amanda D. Melin

## Abstract

Arboreal camera trapping is an emerging remote sensing tool for building plant-frugivore interaction networks with enormous potential to provide new insights into seed dispersal as an eco-evolutionary process. Interaction data derived from camera trap videos broadly falls into two categories: visitation–indicating the presence of an animal in a tree–and intake–indicating true fruit consumption. These linked approximations of seed dispersal may or may not provide distinct sets of information, shaping our understanding of network dynamics, yet the extent to which emergent network structure depends on variable choice is unknown. To address this gap, we deployed camera traps in the canopy of 21 plant species in the Costa Rican tropical dry forest, an endangered biome with sparse community-wide plant-frugivore interaction data. Our objectives were: 1) Quantify plant-frugivore network structure in the tropical dry forest via network, group, and species-level metrics; and 2) Compare metrics between visitation and intake networks. We find strong structural patterns of low connectance, low nestedness, high modularity, and low niche overlap that differ significantly from null expectations in the same direction in both networks. The visitation network exhibited significantly greater modularity & species degree than the intake network, while the intake network exhibited significantly greater plant niche overlap than the visitation network. These findings suggest that the likelihood of consuming fruit on camera differs across species, which may have downstream consequences on other emerging metric estimations. Researchers seeking to build quantitative plant-frugivore networks should therefore consider visitation and intake variables in concert when relying on camera trap data. This work provides foundational plant-frugivore interaction data for a critically endangered biome and enhances our understanding of camera trapping as a remote sensing tool in frugivory and seed dispersal research.

## Introduction

Understanding the eco-evolutionary dynamics between fruiting plants and their seed dispersers is a long-running line of biological inquiry (Eriksson, 2016; Fleming & Kress, 2013). This field of study has been revolutionized by the widespread application of network approaches, which give researchers the ability to model complex webs of species interactions (Lau et al., 2017; Delmas et al., 2019). Plant-frugivore interactions, like other animal-plant systems, are organized into bipartite networks, consisting of two distinct sets of nodes (plants and animals) connected by links (interactions; Carlo & Yang, 2011). To evaluate emergent network structure and detect patterns, ecologists have developed metrics such as nestedness, modularity, connectance, and degree, yielding insights across ecological levels, ranging from the community-to the species-level (Lau et al. 2017).

At present, plant-frugivore network ecology is a burgeoning area of investigation with implications spanning broadscale ecosystem function to species-level adaptation (Acevedo-Quintero et al., 2020; Rumeu et al., 2020). Detailed interaction data have now been generated for a wealth of ecosystems across the world (Gautier-Hion et al. 1985; Palacio et al. 2016; Carreira et al. 2020; Quintero et al. 2020; Bender et al. 2021; Villalva et al. 2024). However, community-wide plant–frugivore network data remain geographically uneven, and Mesoamerican tropical dry forest–one of the world’s most endangered terrestrial biomes– remains comparatively understudied (Almeida & Mikich 2018; Ramos-Robles et al. 2018). This limitation is consequential: this ecoregion is a biodiversity hotspot with pronounced seasonality and strong variation in fruit availability, which may generate interaction structures and species roles that differ from other biomes (Gillespie et al. 2000). Expanding interaction data from Mesoamerican tropical dry forests will therefore improve our understanding of the sources of global variation in plant–frugivore networks while also clarifying the ecological roles of frugivores in these endangered ecosystems.

Furthermore, methods for quantifying plant-frugivore interactions are variable, and include direct in-person observation of fruiting trees, transect surveys, mist-netting for birds and bats, and DNA metabarcoding from fecal samples (Schlautmann et al., 2021; Vitorino et al., 2022) Each of these methods has benefits and limitations, reviewed in Quintero et al. (2022). Among these, arboreal camera trapping is a novel remote sensing tool that is gaining popularity for monitoring plant-frugivore interactions (Quintero et al., 2022; Villalva et al., 2024; Zhu et al., 2022). Camera traps are typically placed in trees directly in front of clusters of fruit, with the cameras set to record 10-s videos upon motion activation (Zhu et al., 2022). From this phytocentric point of view, the objective is to record animals who interact with the fruit. This approach offers several tangible benefits: it is minimally invasive and remote, allowing 24/7 recording of frugivore activity. This is in contrast to direct in-person observation, which can repel cryptic animals, is extremely time intensive, and poses visibility challenges in dense forests with tall canopies (Zhu et al., 2021). Further, camera traps can capture difficult-to-observe species, like bats and other cryptic nocturnal mammals, allowing a fuller characterization of the frugivore assemblage (DePasquale et al., 2025).

While advantageous in several ways, monitoring fruit consumption via camera traps also presents unique challenges. Chiefly, camera trap videos often end before a fruit consumption event is recorded, leaving some outcomes uncertain. This results in two distinct types of possible camera trap data: visitations (i.e., the presence of an animal in a tree) and intake (i.e., direct feeding observations). For researchers seeking to build a quantitative ecological network using camera trap data, the choice of which variable to use will depend on the question at hand, requiring careful consideration of the tradeoffs between visitation and intake networks. For instance, visitation networks are inherently larger, which may allow for a more comprehensive evaluation of community-wide interactions, but may include several non-mutualistic partners, including seed predators, pulp specialists, and animals that occupy trees without feeding. Intake networks, on the other hand, capture a stricter measure of plant-frugivore mutualism, but risk excluding legitimate seed dispersers that are less likely to be captured ingesting/removing fruit on camera (Simmons et al., 2018). Understanding the relationship between visitations and intake is paramount for a nuanced assessment of interaction patterns across a given plant-frugivore community, yet we currently know little about how a given variable (visitation versus intake) impacts emerging network properties across ecological levels, from the network to the species-level.

In this paper, we combine arboreal camera trapping and ecological network analysis to quantify structural patterns of plant-frugivore interactions in a Costa Rican tropical dry forest community. Our objectives are two-fold: 1) Quantify plant-frugivore network structure via network, group, and species-level metrics; and 2) Compare metrics between visitation and intake networks. Ultimately this study will improve both our ecological and methodological understanding of plant-frugivore networks in the critically endangered biome of the Mesoamerican tropical dry forest.

## Methods

### Study system

We studied plant-frugivore networks in Sector Santa Rosa (SSR), Área de Conservación Guanacaste, Costa Rica from January to November 2024. This site is situated in a tropical dry forest in various stages of regeneration following intense restoration efforts beginning in the 1970s (Janzen & Hallwachs, 2020). SSR experiences two distinct seasons: a hot dry season from January to May, and a cooler rainy season from May to December. Habitat-wide ripe fruit abundance is also highly seasonal at this site, with a peak in April during our 2024 field season (Fig. 1). Santa Rosa hosts a diverse and well-documented community of frugivores, including up to 16 species of frugivorous bats, 13 families of frugivorous birds, 3 species of primates, and several other families of mammals (i.e., Fleming, 1979; Janzen & Martin, 1982; Hirsch, 2010) as well as a highly diverse plant community, estimated to host up to 700 vascular plant species across the landscape (Janzen 1988).

**Fig 1.**
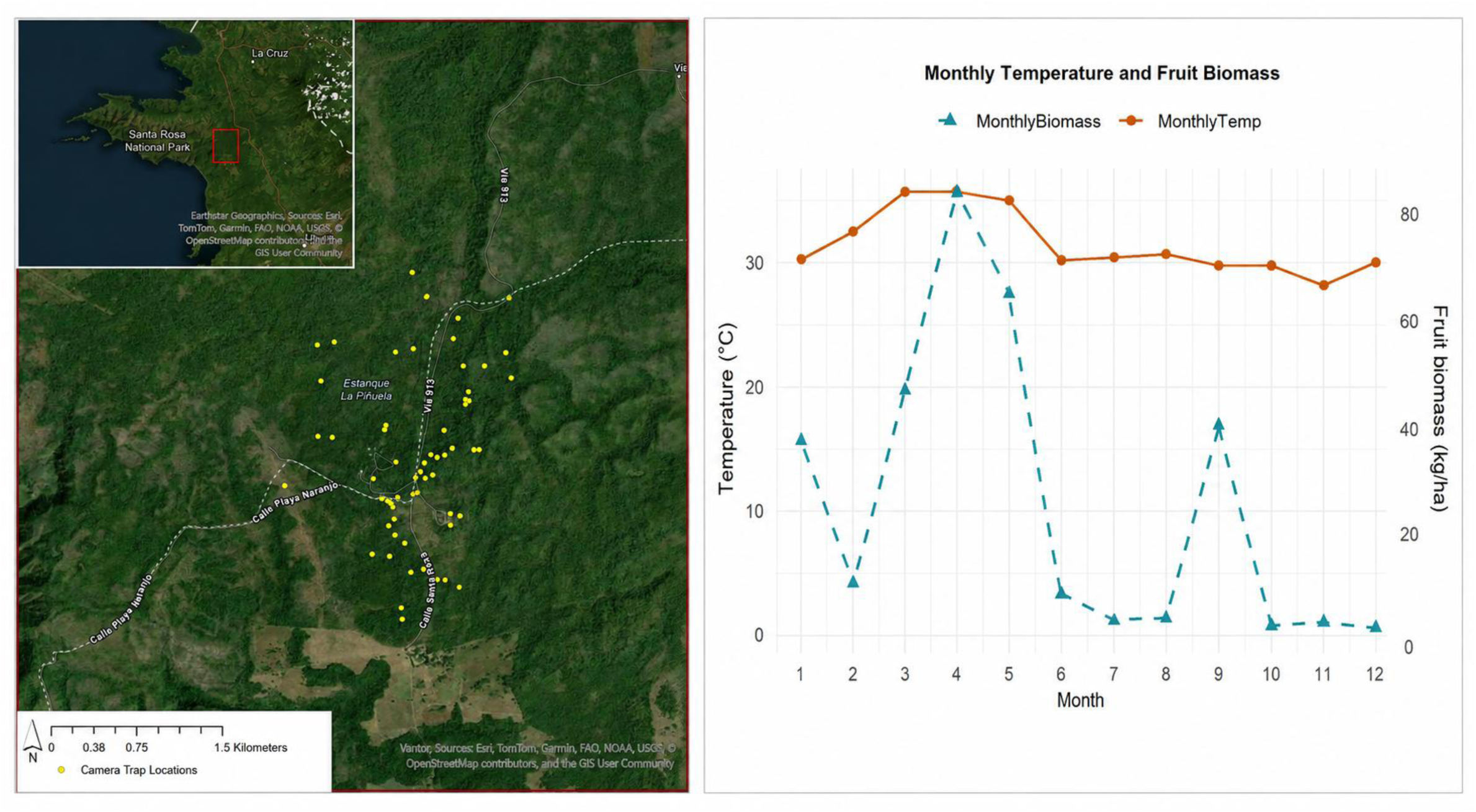
Camera trap locations, monthly temperature, and habitat-wide ripe fruit biomass for the 2024 study period.

### Data collection

We deployed arboreal camera traps (1 per tree) in 66 individual trees across 21 plant species, aiming for 3-5 individuals per species (Figure 1; Table 1). We selected plant species to represent the range of fleshy fruit phenotypic diversity present in SSR, including fleshy and dehiscent fruits of various sizes and colors.The number of individuals studied per species varied based on phenology (i.e., individuals of some species failed to produce fruit), as well as occasional constraints in available equipment (e.g., if a camera was undergoing repair). Cameras were deployed to capture the duration of ripe fruit availability in a given tree individual, which varied from tree to tree and species to species but generally lasted 2-3 weeks. At each tree, cameras were deployed facing ripe fruit within 1-5m. Camera angles were chosen to maximize ripe fruit in the camera’s field of view, but varied depending on individual tree structure (Zhu et al. 2021; Zhu et al. 2022). Cameras were generally set to record 10-sec videos with a 2 minute delay between captures, although we would occasionally increase the video length for larger fruit species (e.g., *Genipa americana*) or ones that require more processing (i.e., *Sloanea terniflora*). Detailed information on sampling effort is given in Table S1, and information on interindividual variability in detections across species is provided in Fig S1.

**Table 1.**
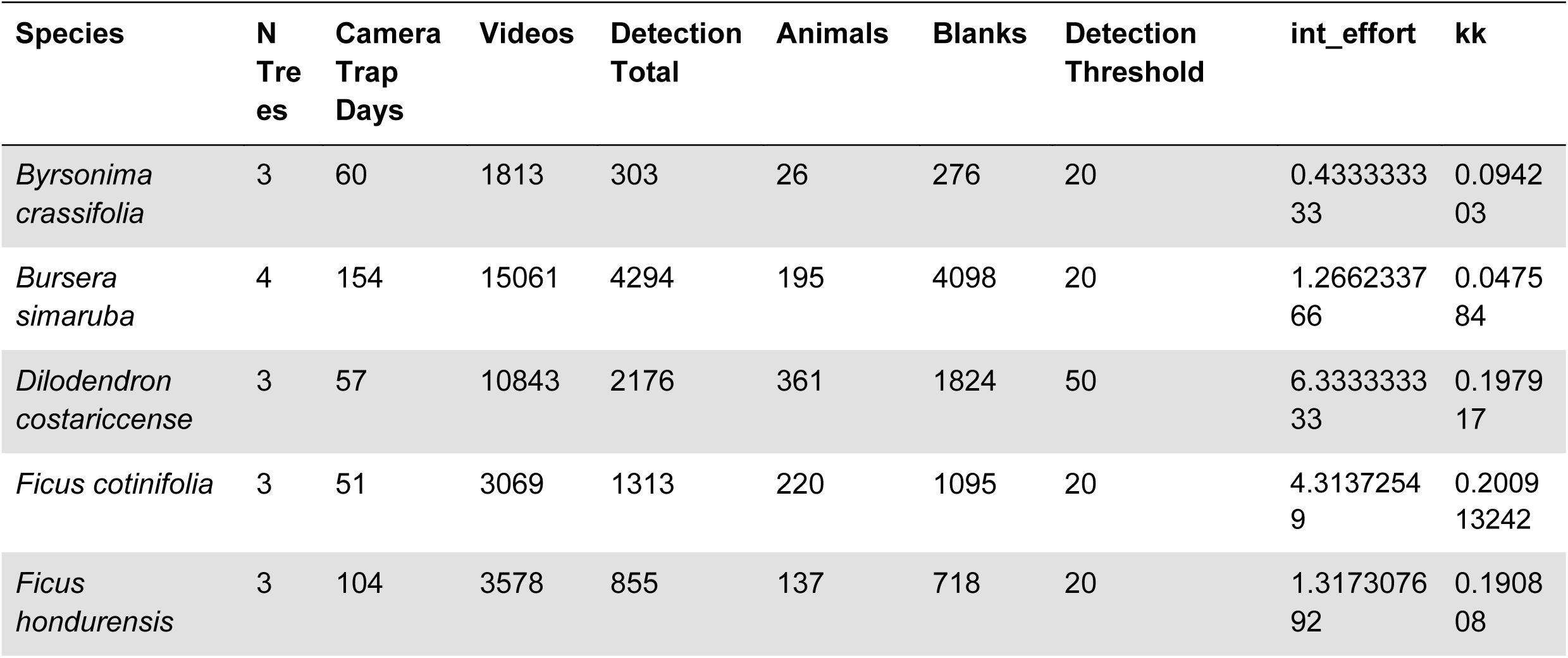
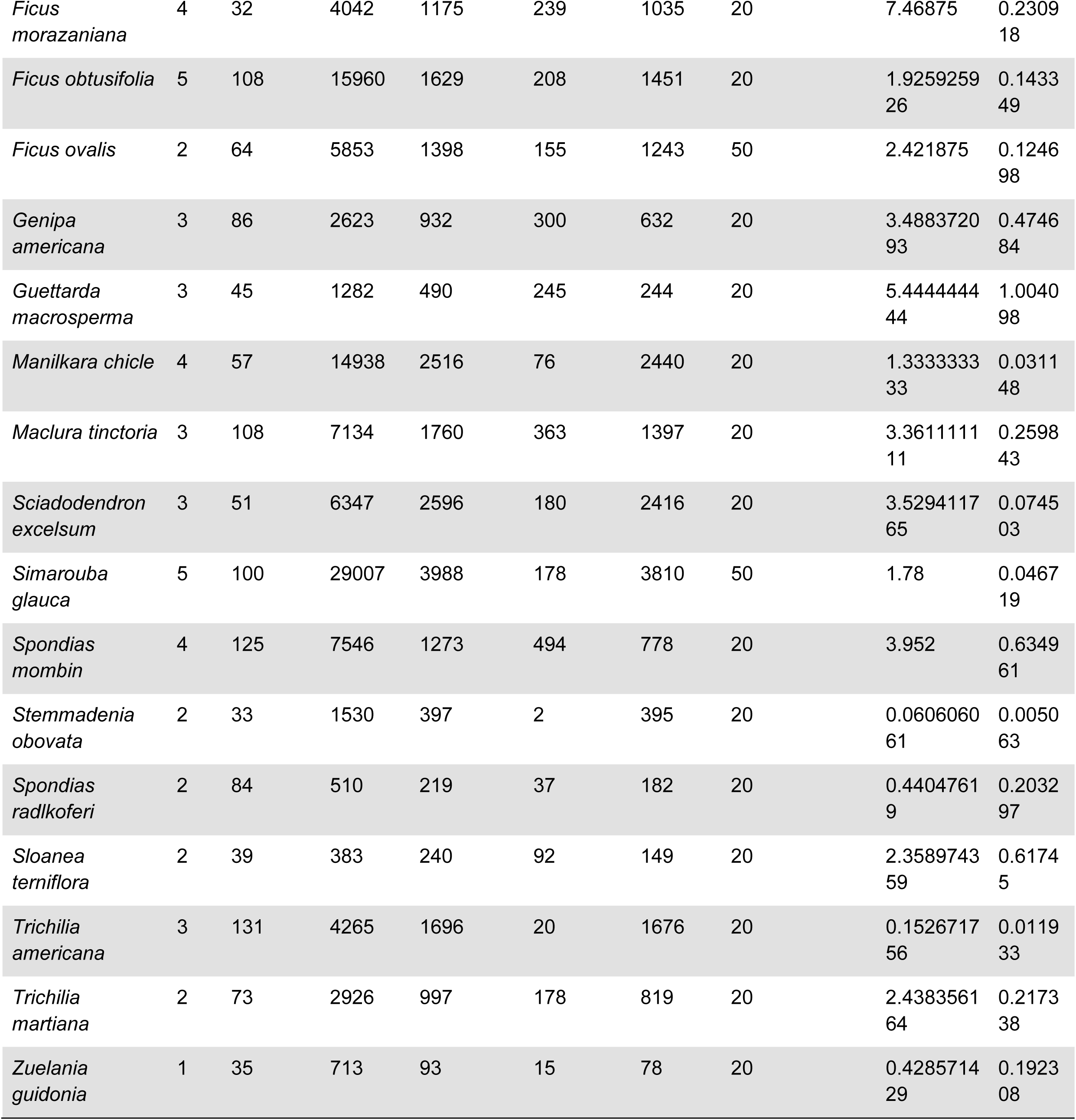
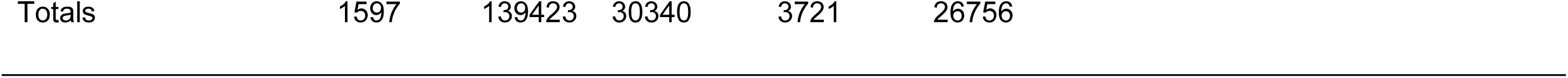
Summary of sampling effort across plant species.

### Data processing

Arboreal camera trapping generates thousands of images, including a majority of false triggers due to wind. The present arboreal dataset yielded a total of 139,423 videos over 1,597 camera trap days, of which ca. 97% were identified as false triggers due to wind. To efficiently process this massive dataset, we used a human-in-the-loop AI image processing approach combining Microsoft MegaDetector and the software Timelapse (Greenberg et al. 2019; Greenberg, 2020; Villalva et al., 2024). This approach uses the MegaDetector algorithm to detect animals, allowing researchers to filter out blank videos based on a confidence threshold, which the researcher can optimize for different tree species and individuals. Detailed information on the detection thresholds used and the sample sizes across tree individuals is given in Table S1. After data processing, the dataset consisted of 3,721 arboreal camera trap videos containing animals (Fig 2). We then coded each animal video, recording: animal species, plant species, tree ID, number of conspecifics present, and number of fruits ingested/removed, based on the outcome seen in the video. We counted a fruit as ingested when it was seen swallowed (i.e., strictest sense of endozoochory), but also included fruits that were deliberately removed by the animal in a foraging context but were not explicitly seen swallowed. Some animals could not be identified to the species level and instead were binned into functionally relevant groupings (i.e., small frugivorous birds, frugivorous bats). Sparse records of known non-frugivores were excluded from the dataset for analyses; these were: anteater (n=1), hawk (n=1), feline sp. (n=2), hummingbird (n=2), nightjar (n=1), vulture (n=1) and owl (n=1), resulting in 3,711 videos in the final dataset.

**Fig 2.**
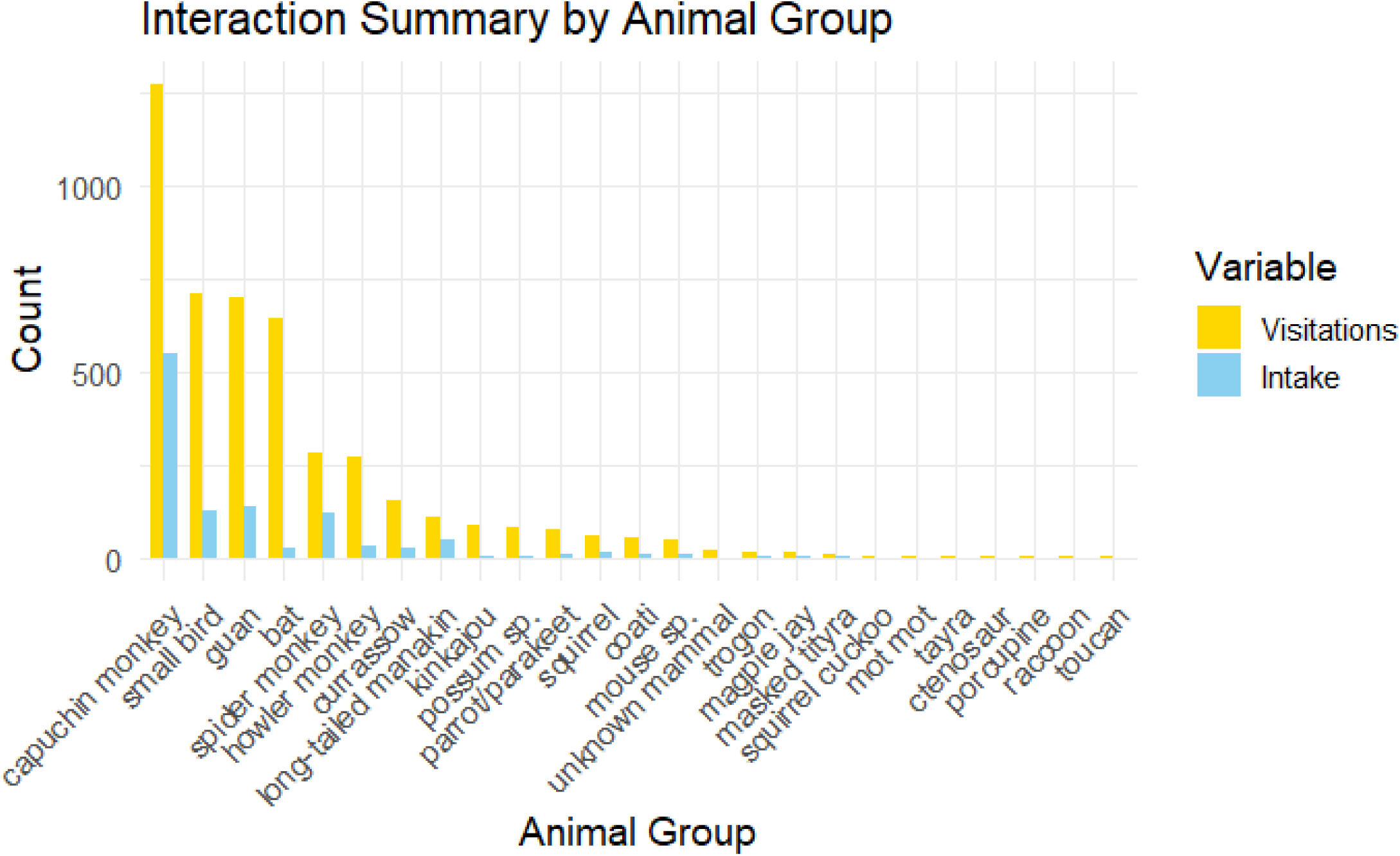
Count of visitation and intake by animal species

### Ecological network analysis

We used R statistical software and the package bipartite (Dormann et al., 2009) to build and analyze plant-frugivore networks based on our camera trap dataset. We built two networks using two different variables from the same dataset: 1) number of conspecifics in each video (a visitation network) and 2) number of fruits eaten/removed in each video (a putative intake network, although the possibility that fruit is dropped outside of the camera frame remains). Networks are visualized in Figure 3. To evaluate sampling completeness across the entire dataset, we constructed accumulation curves of unique-plant frugivore interactions, as well as frugivore species (Figs S2 and S3). For each network, we calculate the following metrics, as defined below:

**Fig 3.**
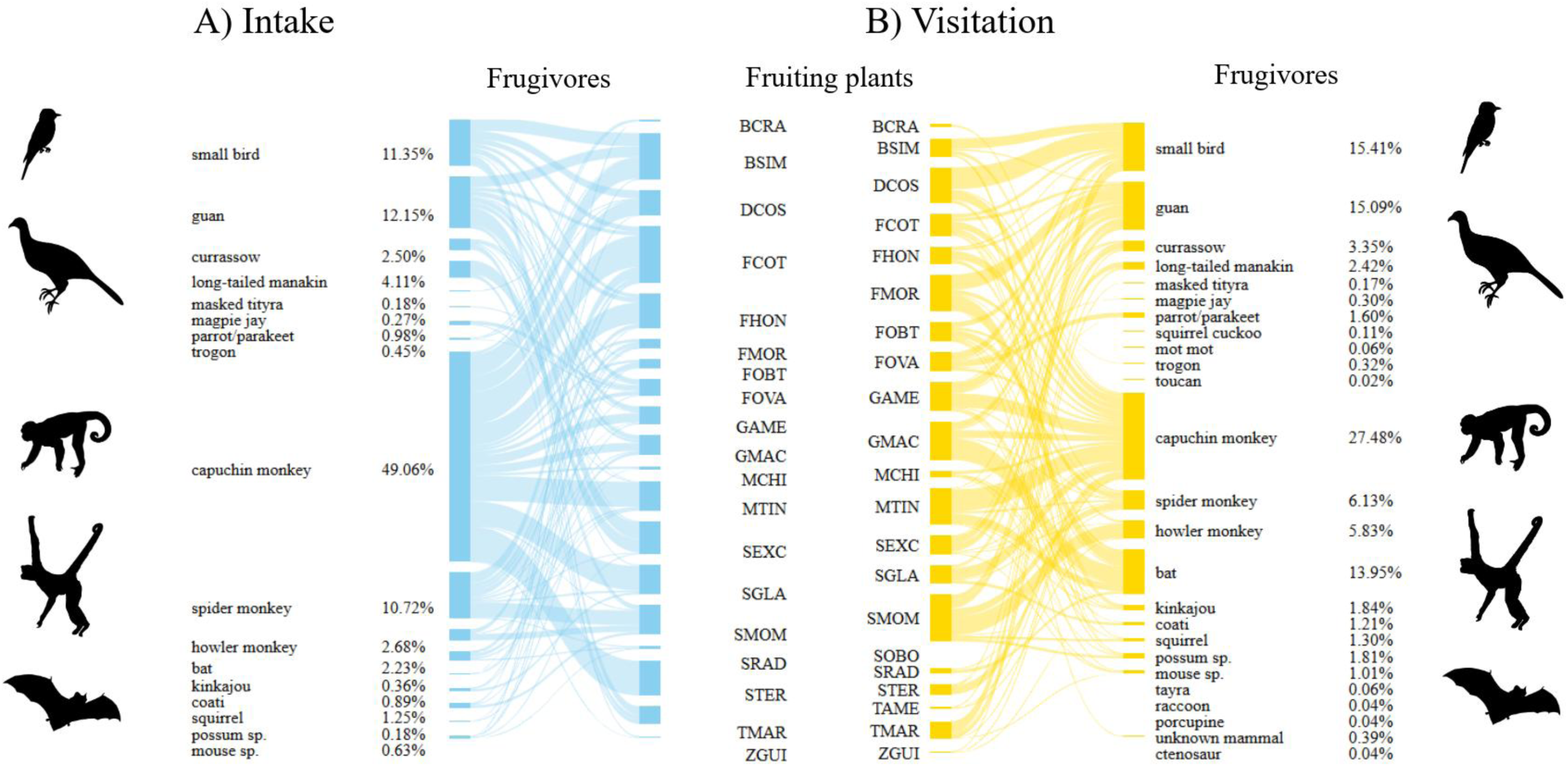
Visualizations of the intake and visitation networks

### Network level

**Connectedness**: Proportion of realized interactions out of all possible interactions

**Nestedness** (via **weighted NODF**): Degree to which specialists interact with subsets of the species that generalists interact with. Higher values indicated a more nested structure.

**Modularity:** Measures degree of compartmentalization in a network, with higher values indicating greater compartmentalization. We employ the compute_modules function from bipartite, set to the default Beckett (2016) community detection algorithm.

### Group level (computes metrics between higher level (animals) and lower level plants)

**Niche overlap**:

Mean similarity in interaction pattern between species of that level, calculated by Horn’s index. Values near 0 indicate no common use of niches, 1 indicates perfect niche overlap.

### Species level

**Species degree**: Sum of links per species.

### Statistical analysis

To statistically evaluate structural patterns in network-and group-level variables, we employ null model analysis, the recommended practice when analyzing ecological network structure (Dormann et al., 2017). This approach allows researchers to determine whether observed network metrics deviate significantly from randomized null expectations, and thus the extent to which observed networks are structured by non-random biological forces, as opposed to neutral processes (Dormann et al., 2009; Neal et al., 2024). In null model analysis, networks are iteratively randomized according to a given algorithm (e.g., randomizing interactions while preserving row and column sums), and network metrics are calculated for each iteration. These “null expectations” can then be plotted alongside the observed network values, and statistically significant deviations calculated using Z-scores.

We generated null models for each network–visitation and intake– using the null_model function in bipartite (Dormann et al. 2008), opting for the default “r2d” algorithm, which randomizes interactions while preserving row and column sums. We then calculated connectedness, nestedness, modularity, and niche overlap for the null models and the observed networks, and plotted both null expectations and observed metrics. Lastly, we calculated Z-scores to evaluate statistically significant deviation from null expectations, and to compare metrics between the visitation and intake networks. To compare our species-level variable (species degree), we generate metrics for both visitation and intake networks. We then conduct a Wilcoxon rank sum test for significant differences between species degrees for the overlapping species between the two networks.

## Results

### Overview of documented plant-frugivore interactions

We recorded 170 unique interactions between 21 plant species and 25 animal species/groups in the tropical dry forest canopy. The visitation network consists of 4,605 interactions (sum of conspecifics per video), while the intake network consists of 1,112 interactions (sum of fruits eaten/removed per video). The breakdown of visitation and intake per animal category is given in Figure 2. Capuchin monkeys (*Cebus imitator*) were the most commonly filmed animal species, present in 30% of videos, followed by unidentified small birds (class Aves; 15% of videos), crested guans (*Penelope purpurascens*; 15%) and frugivorous bats (Order Chiroptera; 8%; see Table 2 for counts and proportions of videos)

**Table 2.**
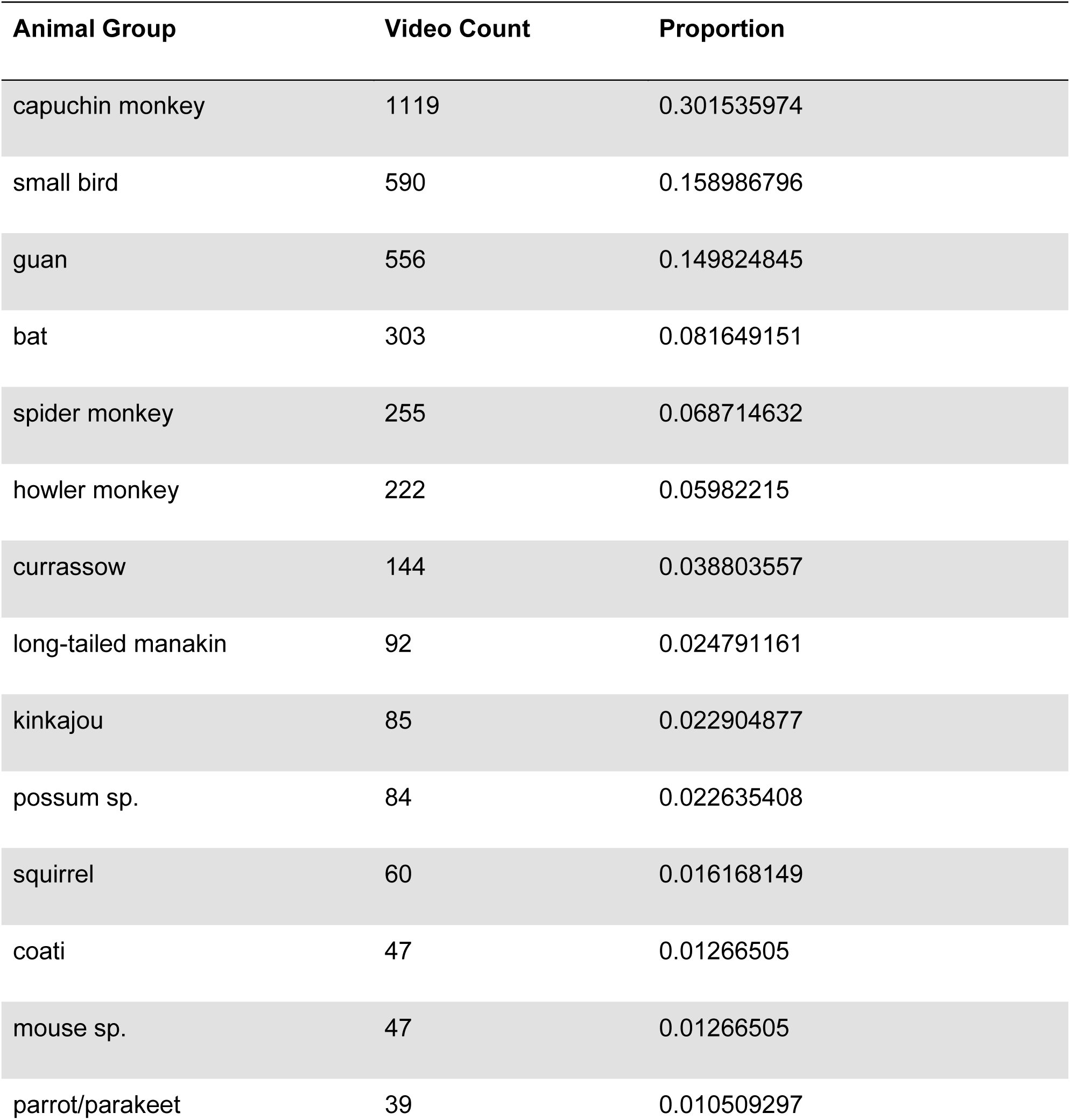
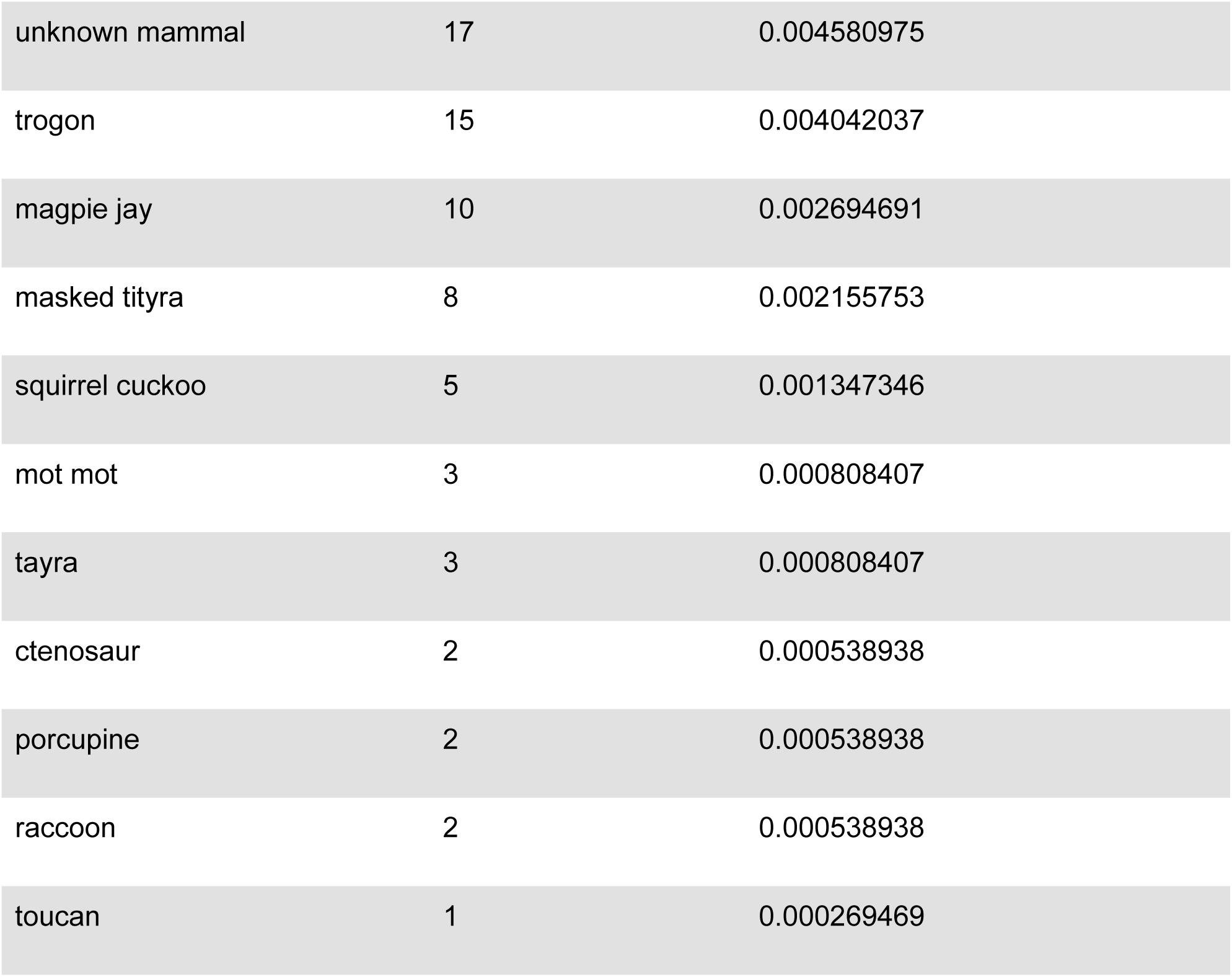
Video count & proportion per frugivore group.

| Animal Group | Video Count | Proportion |
| --- | --- | --- |
| capuchin monkey | 1119 | 0.301535974 |
| small bird | 590 | 0.158986796 |
| guan | 556 | 0.149824845 |
| bat | 303 | 0.081649151 |
| spider monkey | 255 | 0.068714632 |
| howler monkey | 222 | 0.05982215 |
| currassow | 144 | 0.038803557 |
| long-tailed manakin | 92 | 0.024791161 |
| kinkajou | 85 | 0.022904877 |
| possum sp. | 84 | 0.022635408 |
| squirrel | 60 | 0.016168149 |
| coati | 47 | 0.01266505 |
| mouse sp. | 47 | 0.01266505 |
| parrot/parakeet | 39 | 0.010509297 |
| unknown mammal | 17 | 0.004580975 |
| trogon | 15 | 0.004042037 |
| magpie jay | 10 | 0.002694691 |
| masked tityra | 8 | 0.002155753 |
| squirrel cuckoo | 5 | 0.001347346 |
| mot mot | 3 | 0.000808407 |
| tayra | 3 | 0.000808407 |
| ctenosaur | 2 | 0.000538938 |
| porcupine | 2 | 0.000538938 |
| raccoon | 2 | 0.000538938 |
| toucan | 1 | 0.000269469 |

### Network-level metrics

We calculated 3 network-level metrics for both the visitation and the intake networks: connectance, weighted nestedness, and modularity. Observed connectance was significantly lower than expected under the null model in both the visitation network (0.324; z = −31.87, p < 0.05) and the intake network (0.266; z = −20.76, p < 0.05; Figure 4). Although the visitation network was more connected than the intake network, this difference was not significant (p = 0.820). Low connectance indicates that only a small proportion of all possible interactions that could occur, actually occur. When compared to randomized null expectations, significantly lower connectance suggests that the observed interactions are highly selective and non-random.

**Fig 4:**
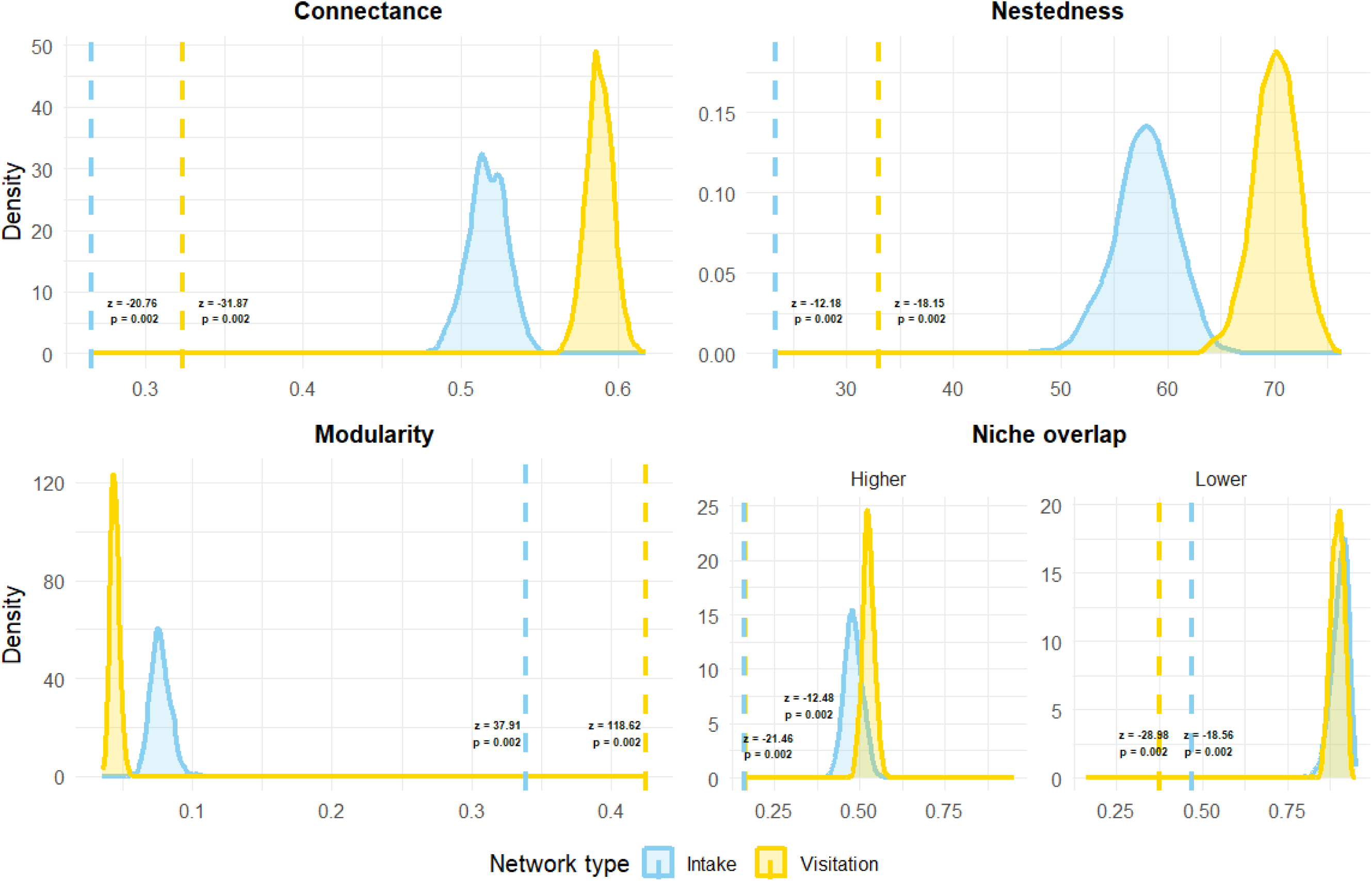
Null distributions (curves) and observed values (dashed lines) for connectance (upper left), nestedness (upper right), modularity (lower left), and niche overlap between animals and plants (lower right).

Likewise, weighted nestedness (WNODF) was significantly lower than null expectations in both the visitation network (33.04; z = −18.15, p < 0.05) and the intake network (23.34; z = −12.18, p < 0.05; Figure 4). The visitation network showed slightly greater nestedness, but the difference between networks was not significant (p = 0.754). Low nestedness indicates that specialists do not interact with proper subsets of the species that generalists interact with, but rather with distinct sets of species.

In contrast to observed connectedness and nestedness values, observed modularity was significantly higher than null expectations for both the visitation network (0.425; z = 118.62, p < 0.05) and the intake network (0.339; z = 37.91, p < 0.05; Figure 4). Modularity was significantly higher in the visitation network than in the intake network (p < 0.05). Modules for both networks are plotted in Figure 5. The nestedness & modularity results together both indicate that the observed networks are characterized by distinct, non-overlapping interaction-dense clusters.

**Fig 5.**
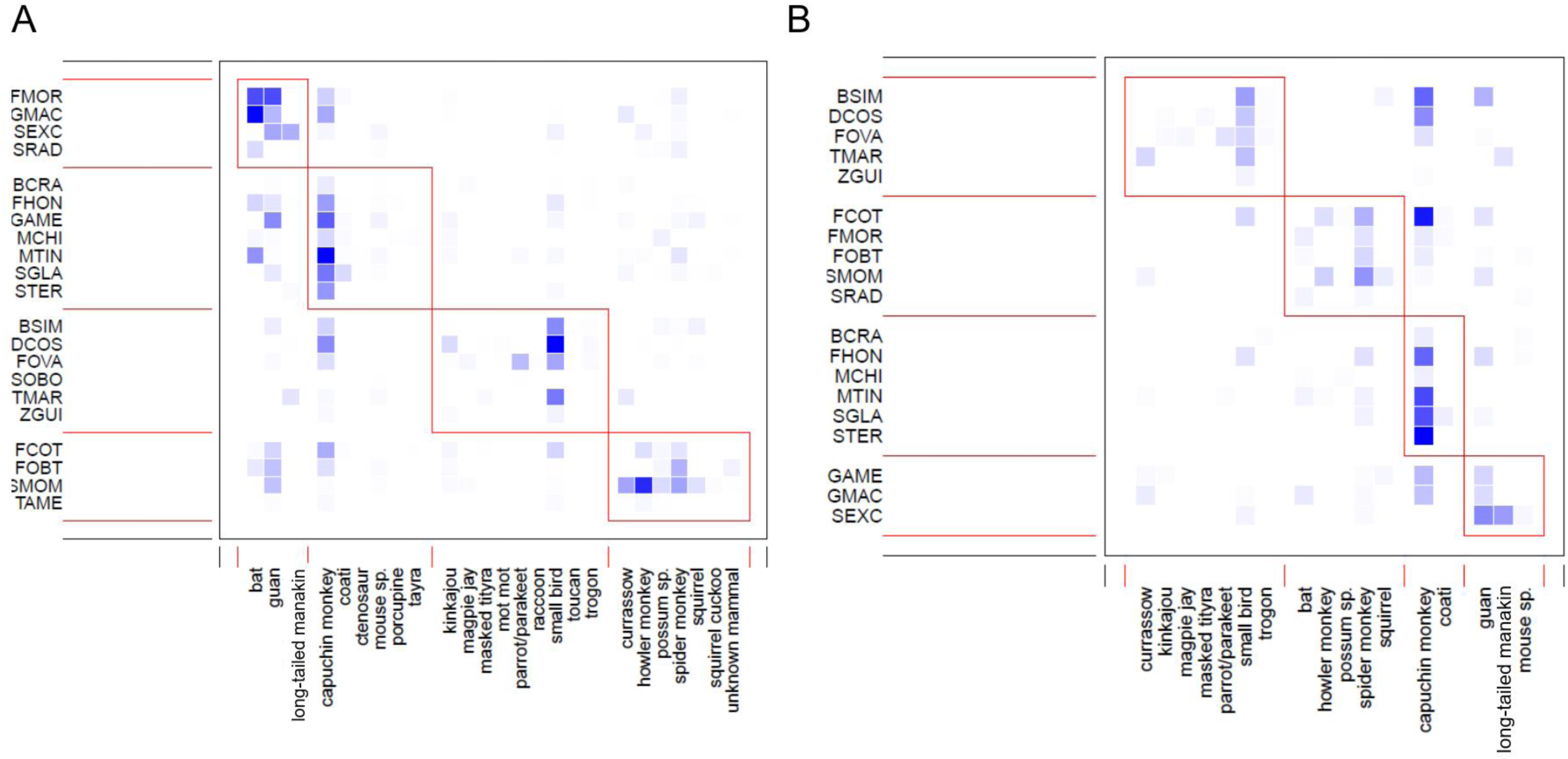
Plotted modules for A) visitation and B) intake networks

### Group-level

We calculated niche overlap (i.e., similarity of interaction patterns) within trophic group-levels: “higher level” denotes frugivores, while “lower level” denotes plants. At both higher and lower trophic levels, observed niche overlap was significantly lower than null expectations. At the animal level, niche overlap was 0.164 in the visitation network (z = −21.46, p < 0.05) and 0.160 in the intake network (z = −12.48, p < 0.001; Figure 4). Observed animal niche overlap was not significantly different between networks (p = 0.973). At the plant level, niche overlap was 0.374 in the visitation network (z= −28.98, p <0.05) and 0.470 in the intake network (z-score= −18.56, p < 0.05), a difference which is statistically significant (p <0.05). These results indicate minimal overlap in resource use between frugivore groups, but moderate overlap in frugivore assemblage between plant species when analyzed using fruit intake.

### Species-level

Species degree differed significantly between networks for both plants and animals, with the visitation network exhibiting higher degree (i.e., more links) than the intake network (Higher level: V = 153, p <0.001; lower level: V = 190, p-value <0.001; Fig. 6). At the animal level, capuchin monkeys had the highest number of links (17 links), while masked tityras (*Tityra semifasciata*) & magpie jays (*Cyanocorax formosus*) had the fewest (1 link). At the plant level, *Spondias mombin* & *Maclura tinctoria* had the highest number of links (13 links), while *Sloanea terniflora* and *Zuelania guidonia* had the fewest (3 links). The extent of pairwise differences in species degree between visitation and intake networks varies across species for both plants and animals.

**Fig 6:**
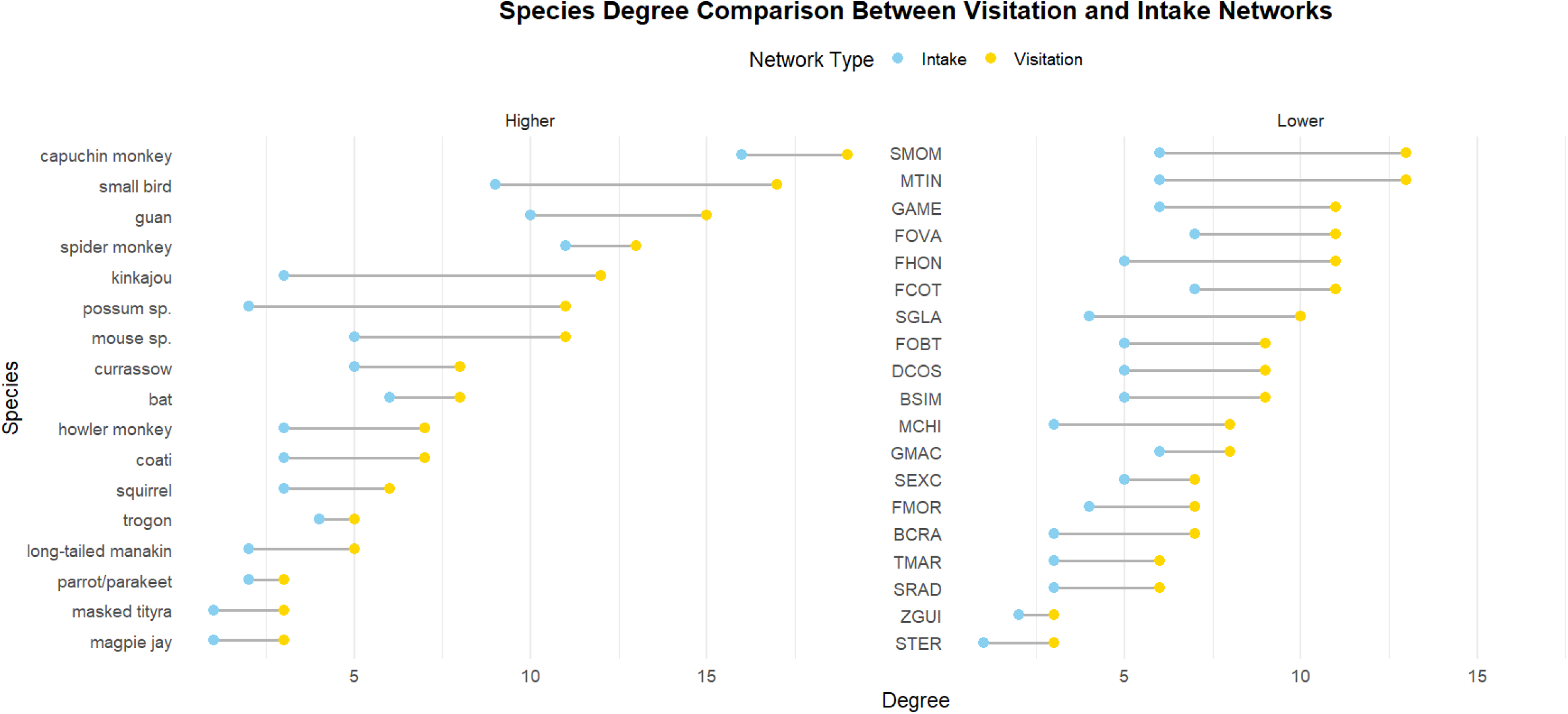
Species degrees between visitation and intake networks

## Discussion

We employed a recently validated method of arboreal camera trapping at fruiting trees to build and analyze two plant-frugivore networks based on two variables common to camera trap video data: visitation and intake. Our goal was to quantify tropical dry forest plant-frugivore network structure and determine whether structural patterns differ between visitation and intake networks. We provide new community-wide interaction data for 21 Costa Rican tropical dry forest plant species. We find that both visitation and intake networks exhibit strong non-random structure, but differ in metrics of modularity, species degree, and plant niche overlap, shedding light on how network metrics vary across ecological levels.

### Objective 1: Quantifying tropical dry forest network structure

Both visitation and intake networks exhibit strong structural patterns, namely low connectance, low nestedness, high modularity, and low niche overlap. This could be due to several biological mechanisms, including strong trait matching between plants and animals, as well as strong niche partitioning between frugivores. This finding is further bolstered by the results for modularity: both networks are more modular than expected under null conditions.

Given the high modularity of the two networks, it is perhaps unsurprising that both networks exhibit significantly lower nestedness, defined as the degree to which specialists interact with subsets of the species generalists interact with. Modularity and nestedness may be inversely related in this system, representing a possible trade off between these two types of network structure (Fortuna et al., 2010). Lastly, niche overlap is significantly lower than null for both plants and animals, suggesting minimal overlap in interaction patterns within animals & within plants. Taken together, these metrics all suggest that the Santa Rosa plant-frugivore community exhibits a strong, non-random modular structure characterized by strong niche separation among frugivores.

Given the diverse nature of the Mesoamerican tropical dry forest system (Janzen 1988; Gillespie and Walter 2001; Powers et al. 2009), these strong structural patterns make sense biologically and shed light on plant-frugivore dynamics. The Santa Rosa community includes several phylogenetically distinct groups of fruiting plants with diverse suites of traits, alongside several distinct groups of frugivores with differing anatomical traits. For example, birds and mammals appear to cluster distinctly in our modularity analyses (Fig. 6). Birds are constrained by gape width, and thus tend to interact with small, dehiscent fruit that they can swallow whole (Lord, 2004; Wheelwright, 1985). Mammals, on the other hand, tend to be larger-bodied and capable of consuming bigger fruit (Goebel et al., 2023). The effect of body size is evident in the modules: only the larger birds– crested guans (*Penelope purpurascens*) and great curassows (*Crax rubra*)–cluster with mammals, likely due to their ability to exploit larger fruits. Interestingly, contrary to what might be expected based on its body size, the nocturnal kinkajou (*Potos flavus*) clusters with the smaller birds, due to its frequent consumption of small dehiscent lipid-rich fruit (i.e., *Dilodendron costariccense*, *Bursera simaruba*). Trait-matching may similarly be occurring along traits such as color, odor, and hardness, given differing capacities for color vision, olfaction, and tactile inspection of fruits (Valenta & Nevo, 2020), which remains to be thoroughly investigated in this system.

### Objective 2: Comparing visitation and intake network metrics

Connectance, nestedness, and niche overlap indices were largely congruent between visitation and intake networks, although plant niche overlap was significantly higher in the intake network than the visitation network. At the network level, only modularity differed significantly between visitation and intake networks, with the larger visitation network being significantly more modular than the smaller intake network. Both networks exhibit four modules, but the exact species within each module differ, suggesting that module assignment is sensitive to the difference between visitations (i.e. potential interactions) and intake (i.e., realized interactions). Some species–due to their inherent characteristics–are less likely to be captured removing fruit on camera, and thus will not appear in the intake network, though they still may be consuming fruits at the trees they visit. Kinkajous and other nocturnal mammals, for example, are rarer to capture feeding directly in front of the camera, but can often be seen engaging in obvious fruit investigation behaviors. Other animals, like crested guans, will “loiter” in fruiting trees by sitting in the same spot without feeding, resulting in a much larger interaction strength in the visitation network versus the intake network. As a direct consequence of species-level traits favoring fruit consumption on camera, estimates of plant niche overlap are significantly higher when estimated using intake versus visitation.

Like modularity, species degree also differed significantly between the visitation and intake network for both plants and animals, suggesting that species-level metrics may also be sensitive to these different variables. Certain species/groups exhibit larger pairwise differences in degree than others. For example, unknown small birds, kinkajous, possums, and mice exhibit larger pairwise differences in degree between visitation and intake networks, suggesting that visitations by these groups are not necessarily reflective of intake, either due to difficulty in capturing feeding events, or due to a biological reality where animals visit trees a given species but do not feed (i.e., as part of travel or other related habitat use). On the other hand, certain groups, like spider monkeys, bats, and trogons exhibit smaller pairwise differences, suggesting that visitation is more tightly linked to intake. This species-level analysis provides insight into these taxon-level differences in visitation and fruit intake, adding to the growing body of work on the role of individual species within networks (Ramos-Robles et al., 2018).

These results suggest that visitation and intake variables should be considered in concert, particularly when animal species identity is concerned, as in metrics of modularity and species degree. Researchers should pay close attention to taxa that cluster differently in each network in order to identify potential species biases. Further, given the broader inclusion of non-mutualistic partners, visitation networks may overestimate modularity when compared to intake networks. For example, some taxa present in the visitation network disappear from the stricter intake network, indicating a discrepancy between visitation and intake for these groups.Ultimately, the choice of which network to use will depend on one’s research questions. Visitation networks would be appropriate for broad community studies that seek to capture the occupancy across the entire assemblage of frugivores, regardless of functional outcome, while intake networks would be more appropriate for studies focused on mutualistic interactions and legitimate seed dispersal.

### Implications

Our study reinforces the utility of arboreal camera trapping for capturing fruiting plant visitations by diverse frugivore groups while highlighting potential pitfalls in their application to build interaction networks. In particular, we note that some animal groups appear more likely than others to consume fruit in front of a camera trap, suggesting that intake networks built using camera trap data may be biased towards certain groups, which should be considered carefully by researchers & practitioners. Capuchin monkeys may be an example of this in our dataset, as they account for half of all fruit intake. While capuchins are known to be abundant and important frugivores in Santa Rosa, their gregarious nature may make them more likely to consume fruit in front of a camera trap. As such, researchers should consider how species traits can shape camera trap outcomes and drive resulting network structure.

### Limitations

The present study is limited by two main factors. First, for the analysis conducted here, we were unable to identify some frugivores to the species-level (i.e., unknown small birds, frugivorous bats, small nocturnal rodents, nocturnal possums). When characterizing a plant-frugivore network, species-level information is ideal for a comprehensive assessment of frugivore species identity. Species-level differences in these binned taxa are likely evident, but we are unable to evaluate them at present. Future work should seek the collaboration of bird, bat, and mammal experts for species identification. Second, due to stochasticity in phenological patterns, we could not always achieve the minimum goal of three individual trees for a given species, resulting in uneven sampling of certain plant species (*Ficus ovalis*, *Sloanea terniflora*, *Zuelania guidonia*). The resulting interactions thus may not be representative of the entire assemblage of frugivores for these species. Further, the community represented here is only a sample of the larger plant-frugivore community in Santa Rosa, which is estimated to contain up to 700 plant species (Janzen 1988). Additional sampling is likely to reveal more interactions.

## Conclusions and future directions

This study represents an important contribution to the field of plant-frugivore network ecology by assessing how network outcomes converge and contrast when using two variables common to camera trap data. We find strong structural patterns in both networks, namely low connectance, low nestedness, high modularity, and low niche overlap. In line with hypotheses of trait-based network assembly, these patterns suggest the presence of strong biological mechanisms operating between plants and frugivores, including trait-matching and niche separation, which also may be shaped by species abundances (Dormann et al., 2017). Further, We find that three metrics–modularity, species degree, and plant niche overlap–differ significantly between the visitation and intake networks. The visitation network exhibited higher modularity and species degree, while the intake network exhibited higher plant niche overlap. Future studies should aim to expand sampling across more individuals and species to further develop our understanding of plant-frugivore dynamics in the tropical dry forest ecosystem. Overall, this study sheds new light on the relationship between visitation and intake in camera trap datasets, and highlights the importance of explicitly analyzing and comparing these variables when characterizing a plant-frugivore community using arboreal camera trapping.

## Acknowledgements

We would like to thank Roger Blanco and the staff of the Área de Conservación Guanacaste for making this work possible. We would additionally like to thank Pablo Villalva for feedback on initial stages of the manuscript and for sharing code to generate the summary table of sampling effort. Funding for this work was provided by the Leakey Foundation (to A.N.D), the American Philosophical Society (to A.N.D), and the Natural Sciences and Engineering Research Council (NSERC) of Canada through the Vanier Canada Graduate Scholarships (to A.N.D.), Discovery Grant (RGPIN-2017-03782 to A.D.M), and Canada Research Chairs (950-231257 to A.D.M.) programs.

---

**Fig S1:**
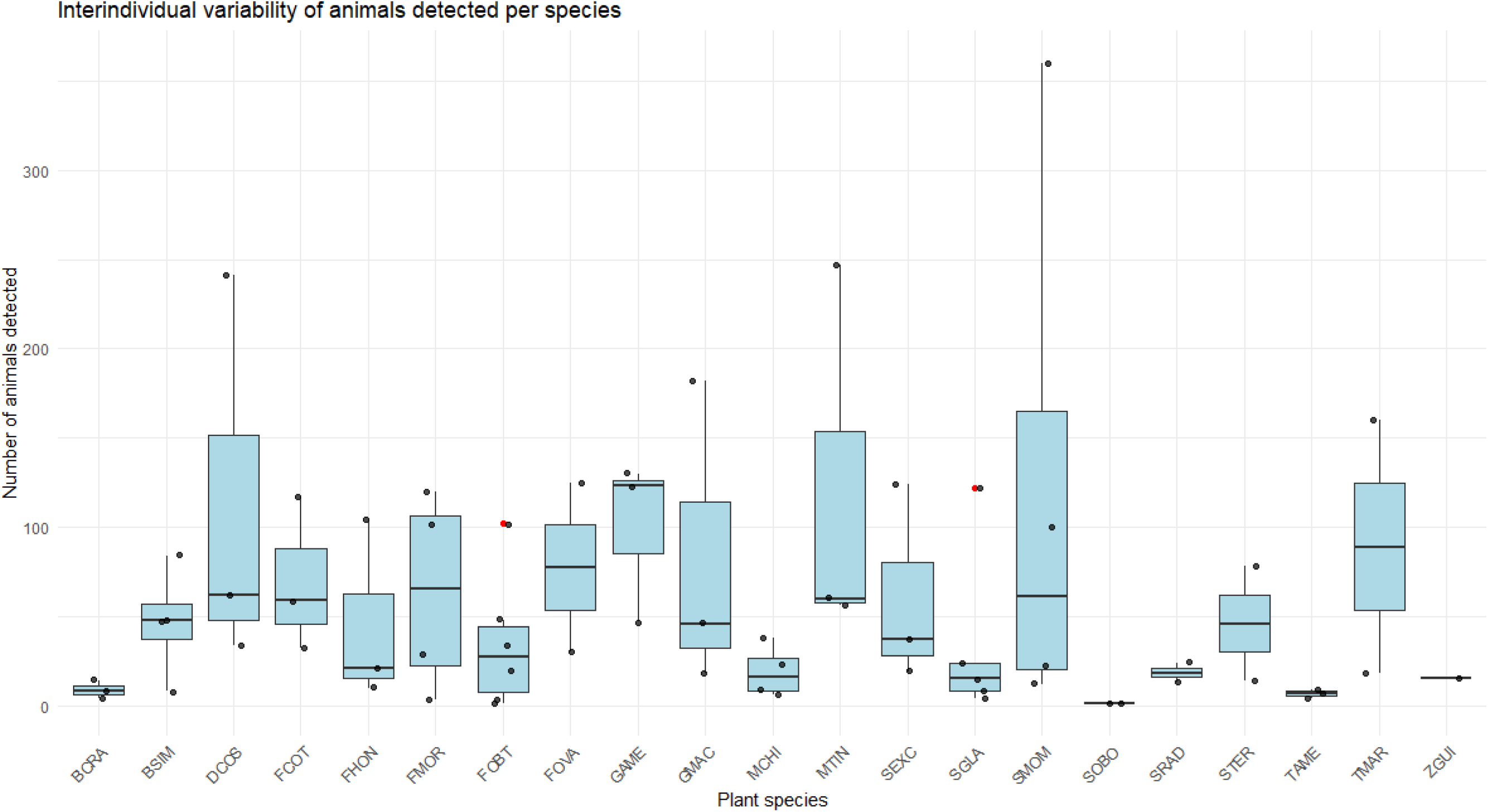

**Fig S2:**
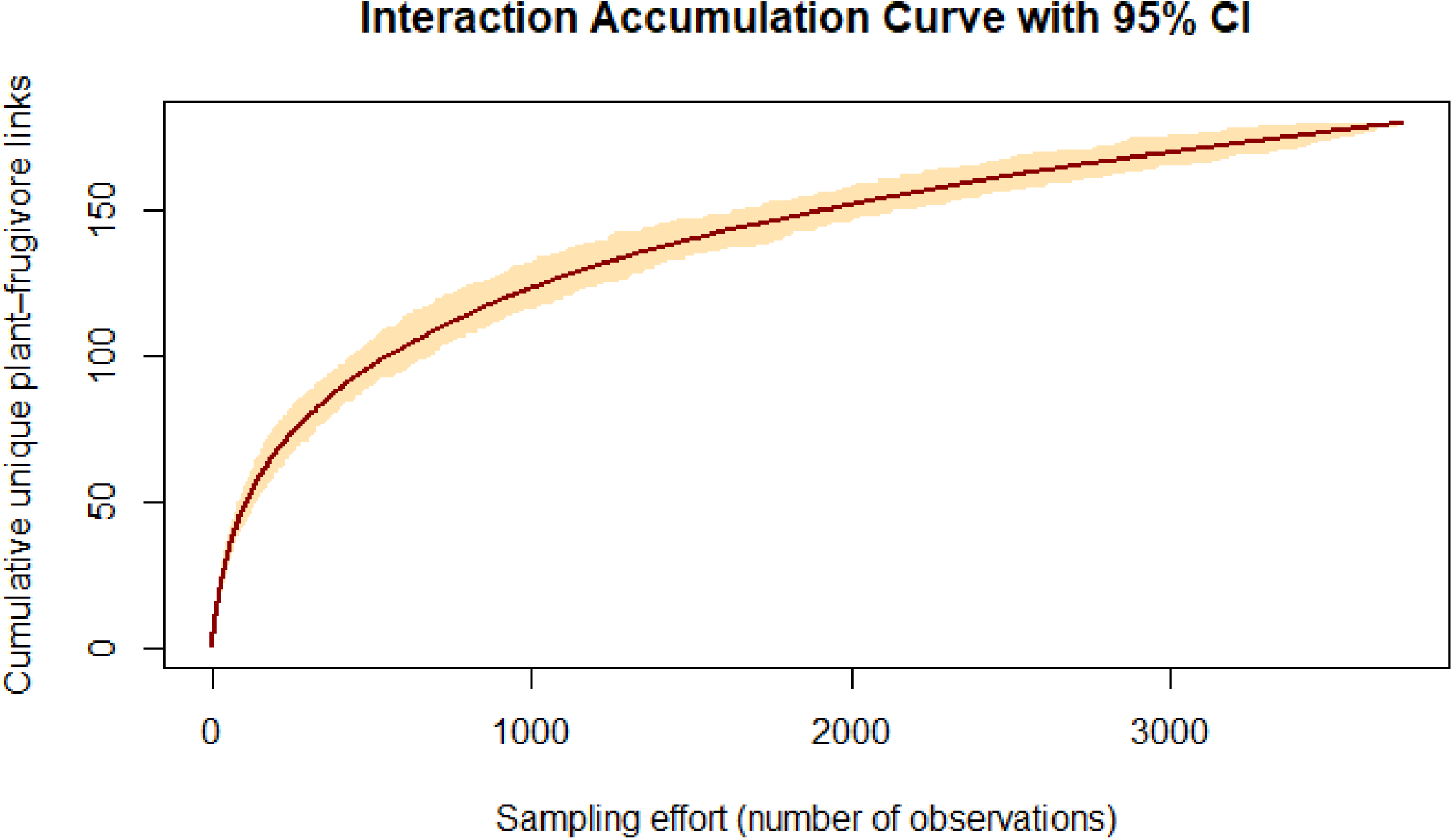
Interaction accumulation curve for unique-plant frugivore links

**Fig S3:**
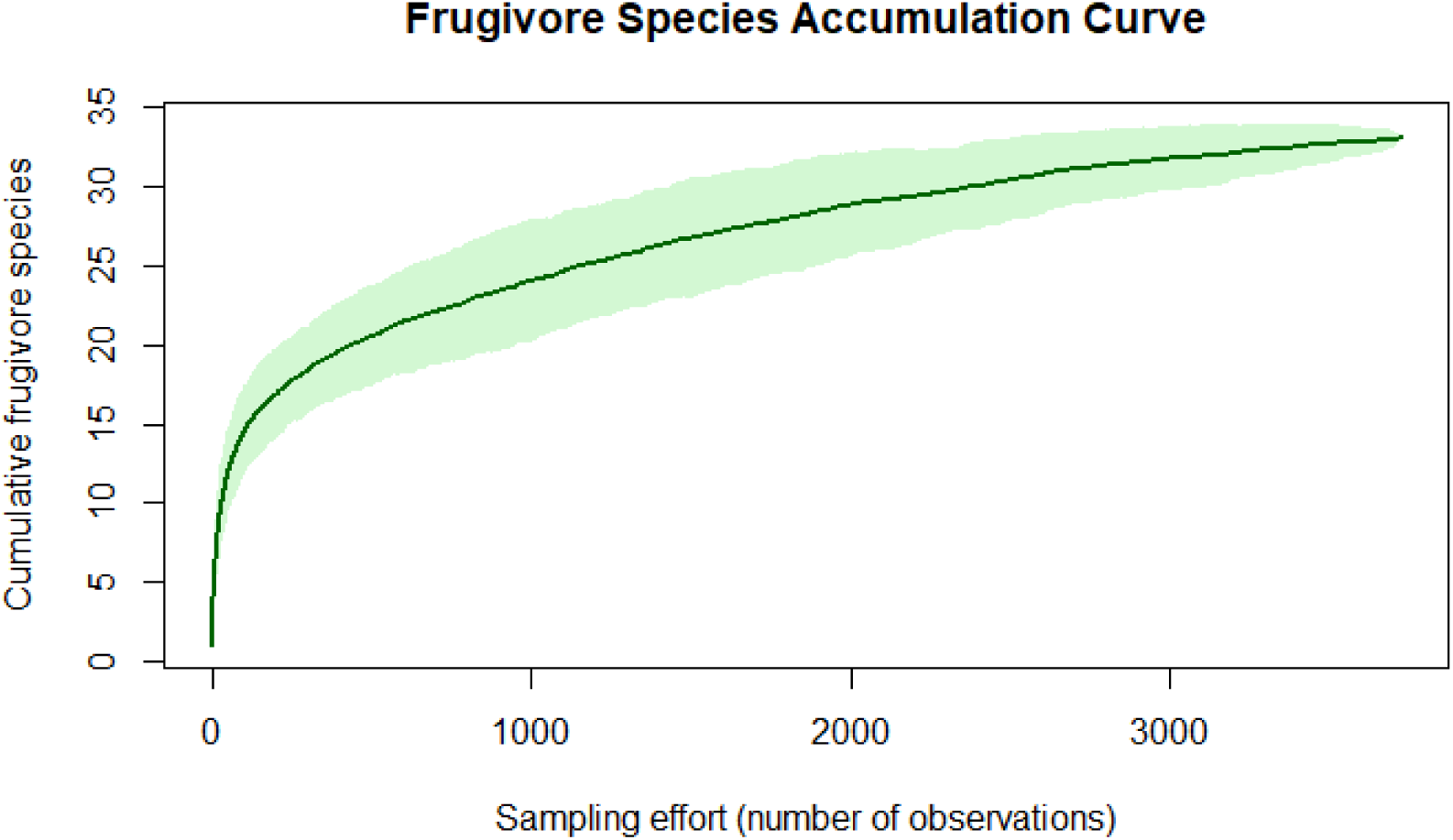
Accumulation curve for frugivore species

**Fig S4.**
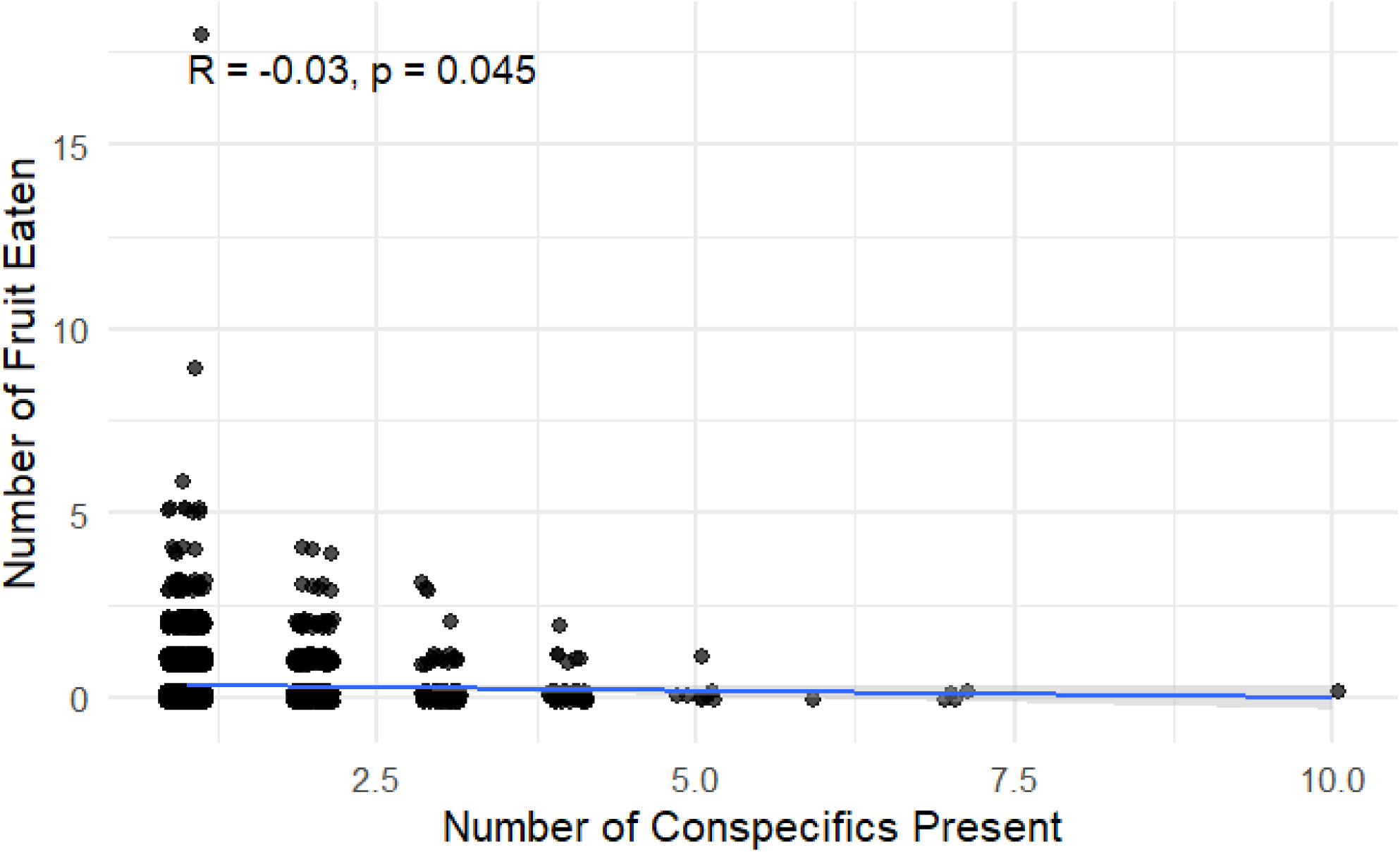
Correlation plot between number of conspecifics present and number of fruit removed per video

**Table S1:**
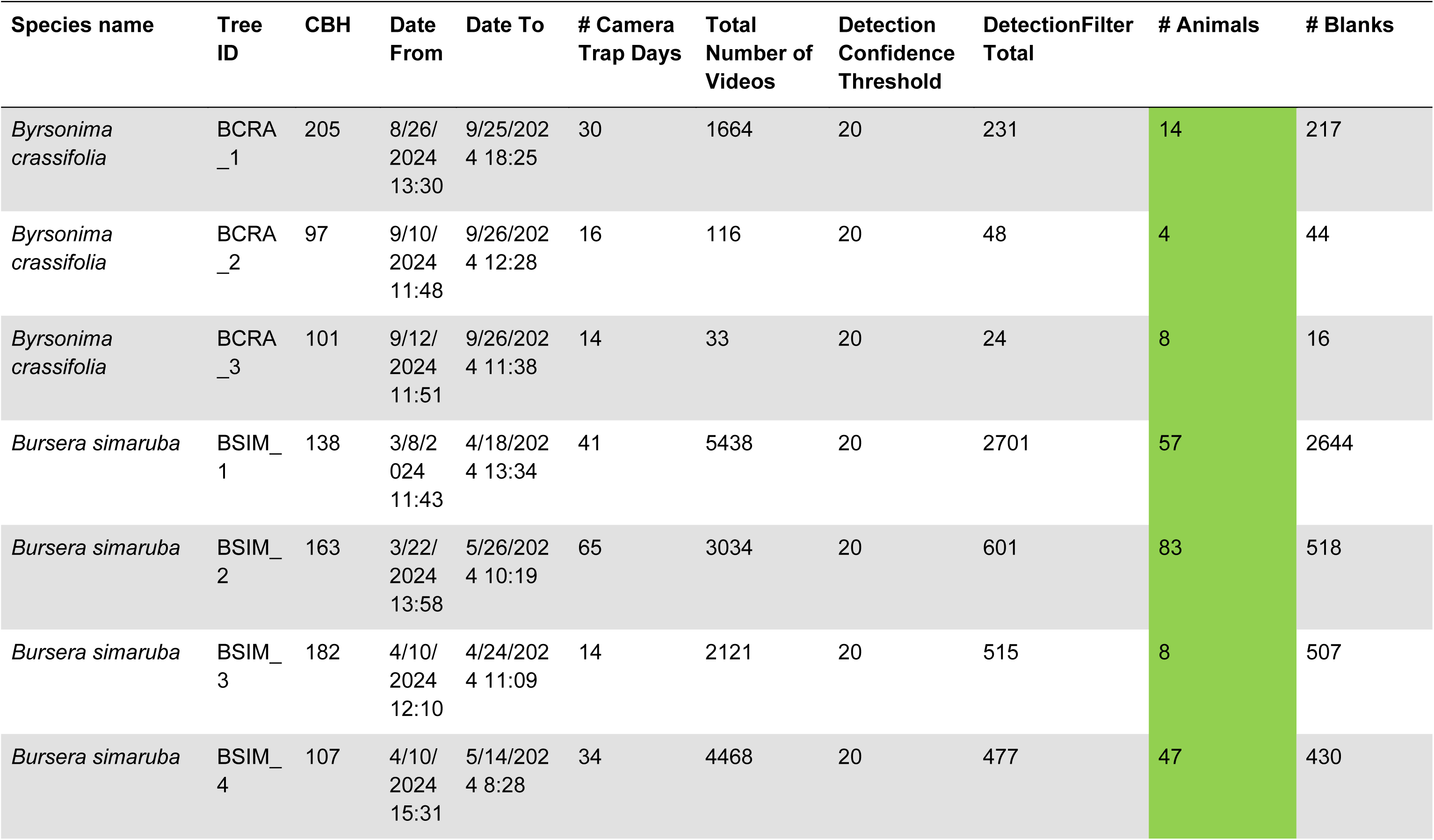
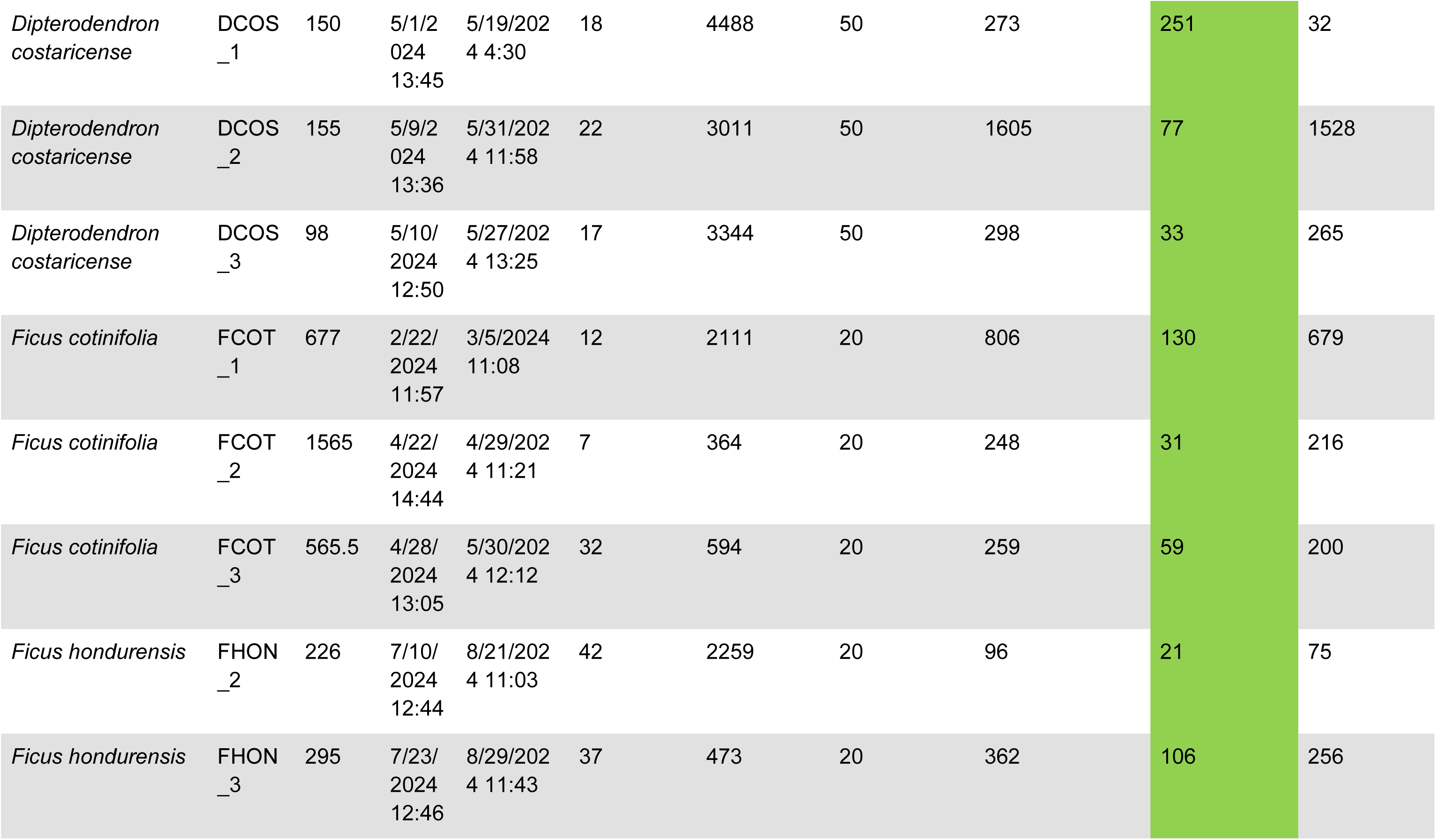
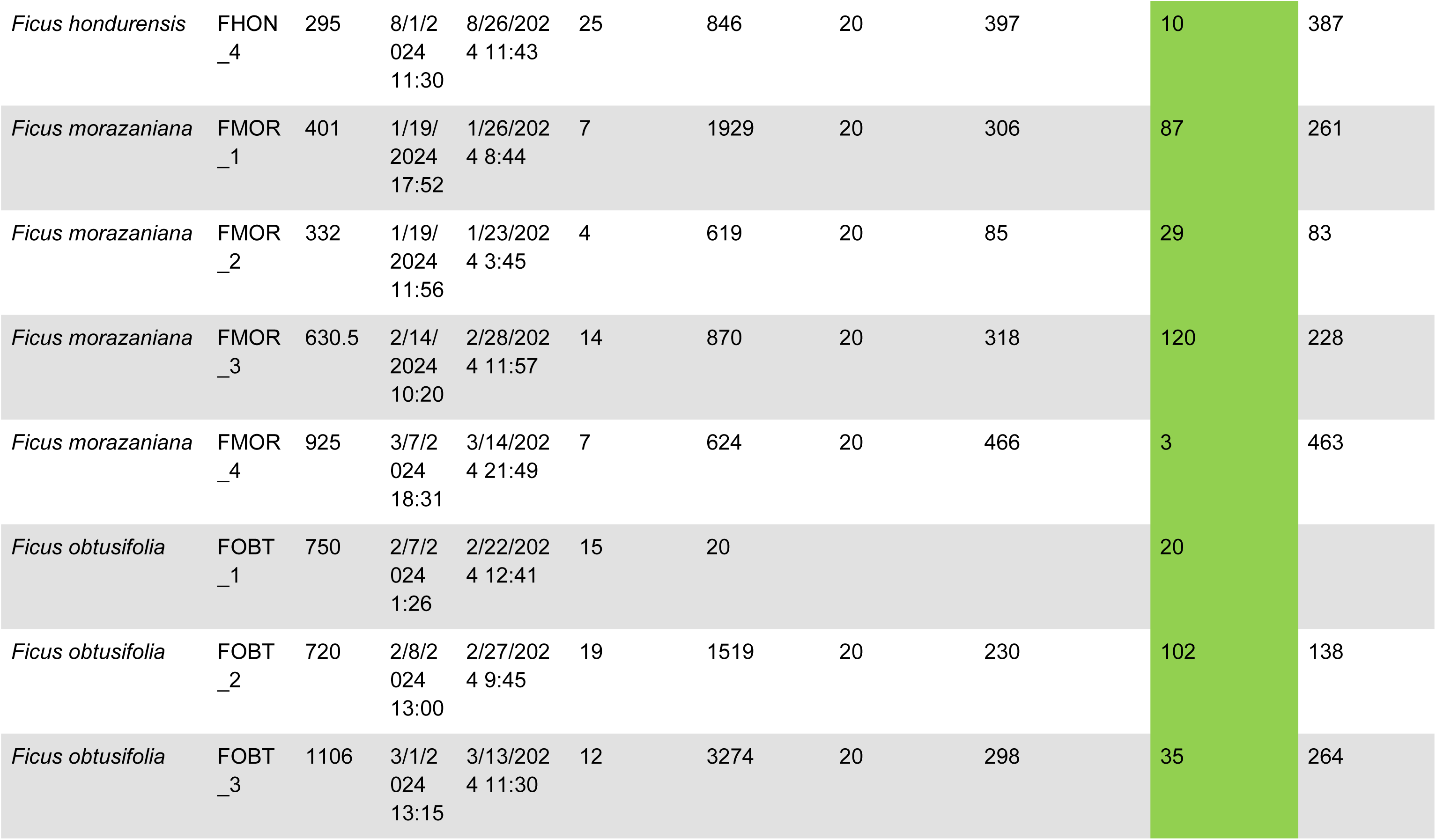
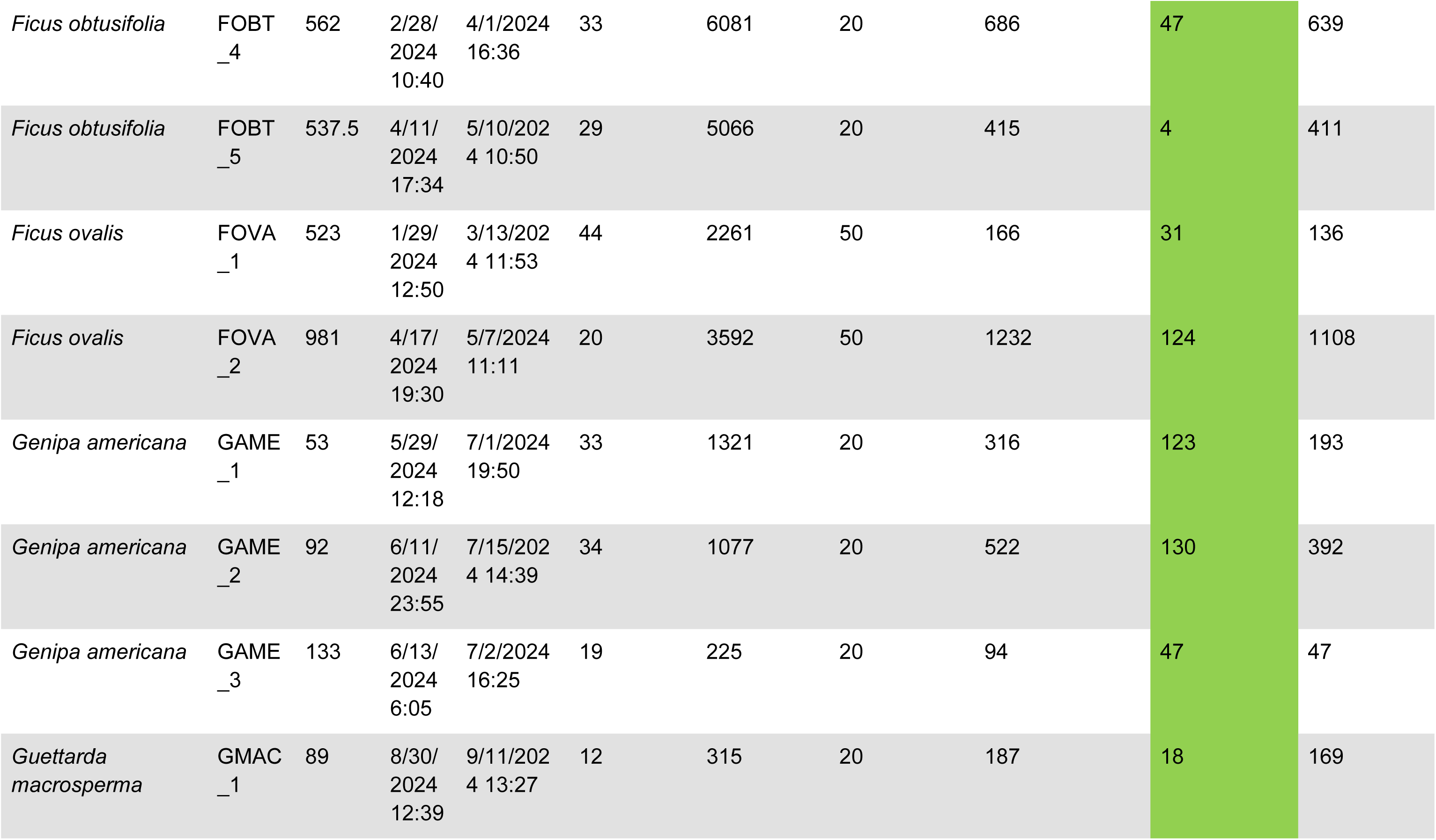
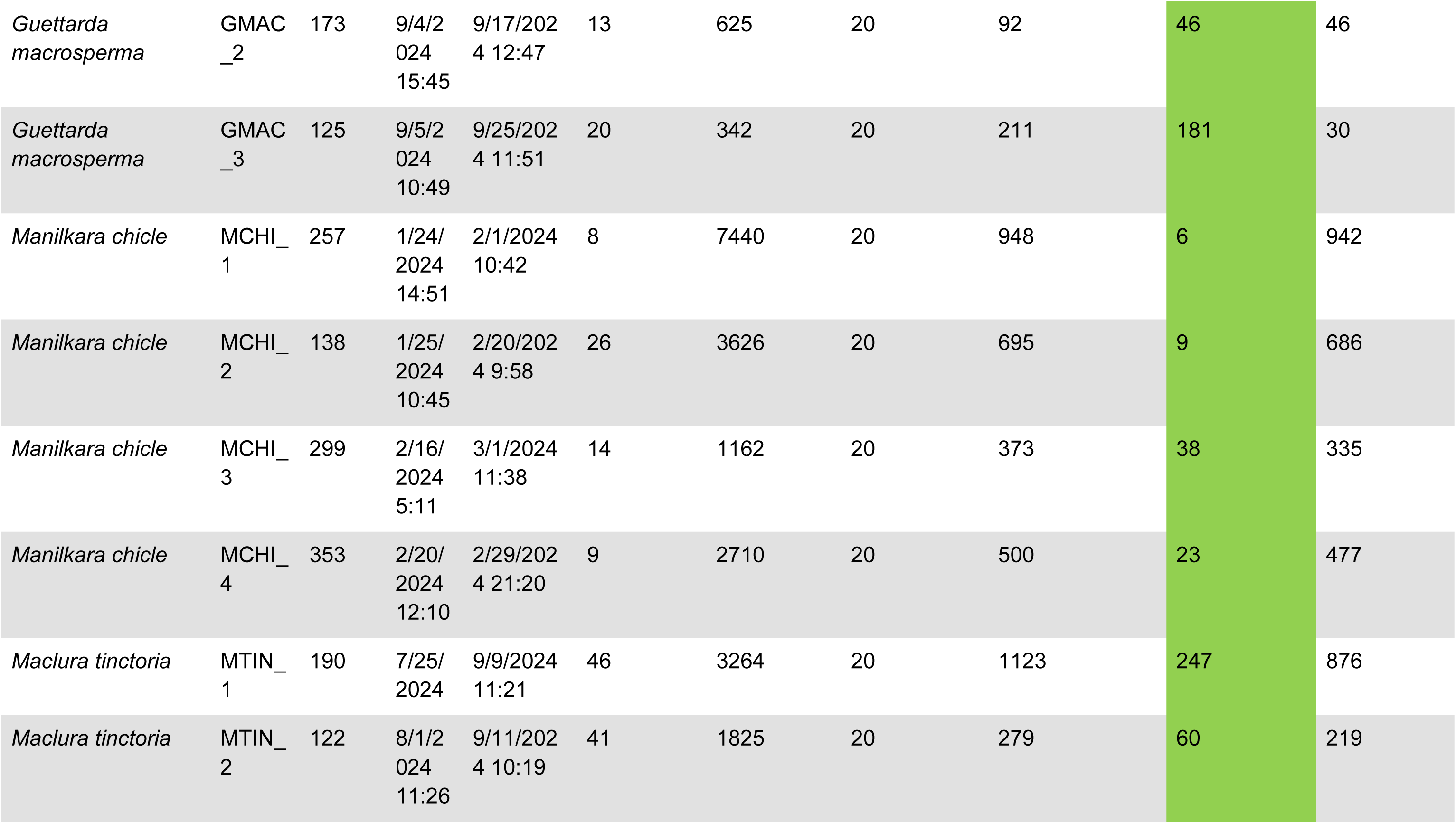
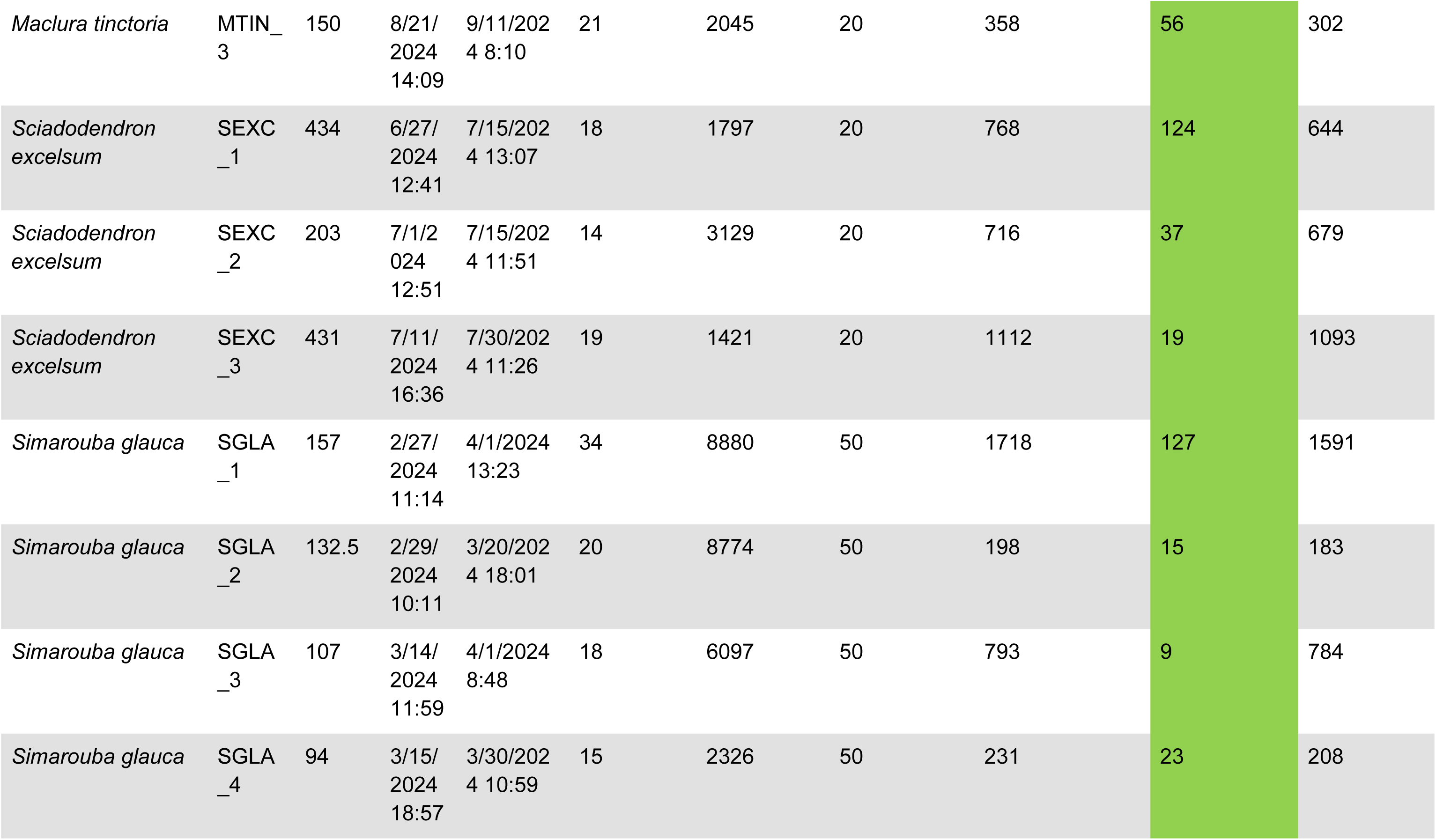
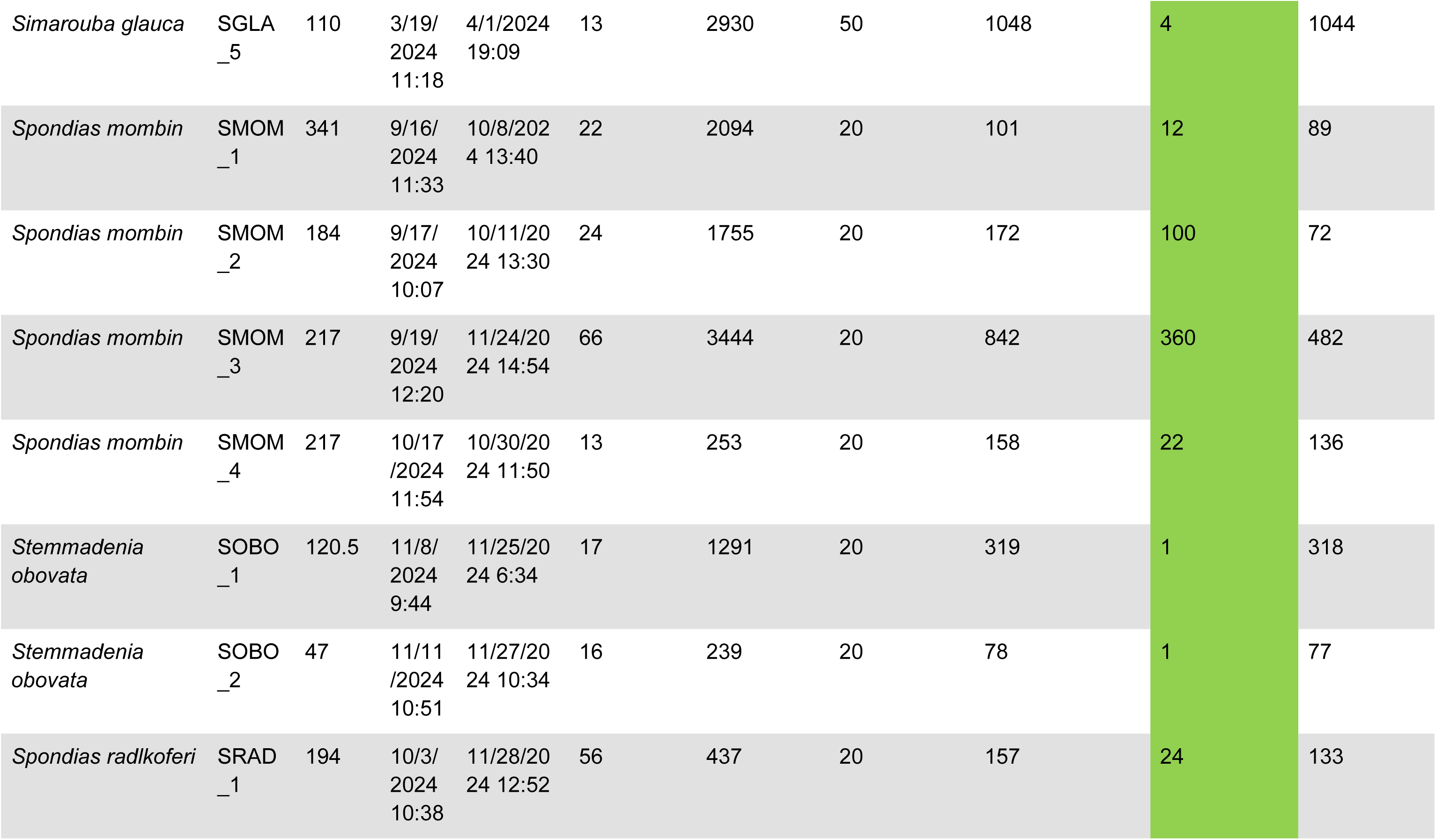
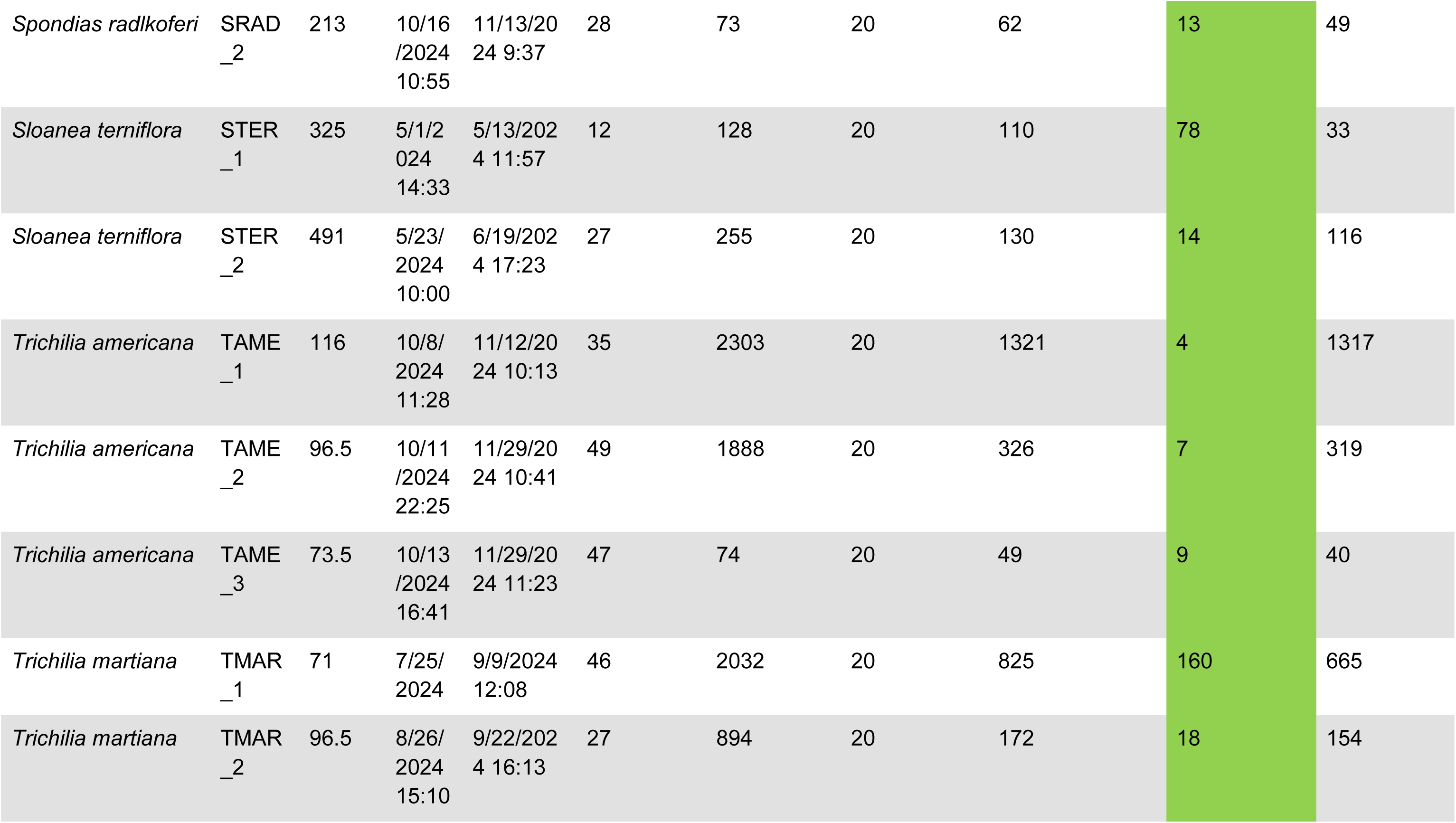
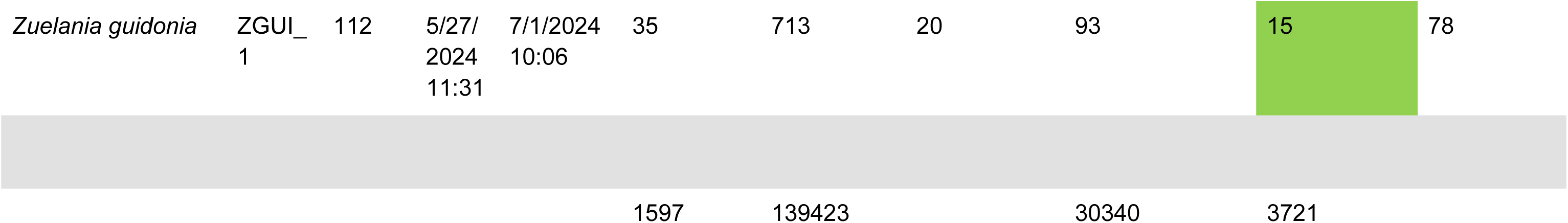
Sampling information by tree individual.

